# Mapping the subcortical connectome using in vivo diffusion MRI: feasibility and reliability

**DOI:** 10.1101/2022.03.28.485689

**Authors:** Jason Kai, Ali R Khan, Roy AM Haast, Jonathan C Lau

**Author notes:** Corresponding author Jonathan C. Lau, M.D., Ph.D., FRCSC, Assistant Professor.

## Abstract

Tractography combined with regions of interest (ROIs) has been used to non-invasively study the structural connectivity of the cortex as well as to assess the reliability of these connections. However, the subcortical connectome (subcortex to subcortex) has not been comprehensively examined, in part due to the difficulty of performing tractography in this complex and compact region. In this study, we performed an *in vivo* investigation using tractography to assess the feasibility and reliability of mapping known connections between structures of the subcortex using the test-retest dataset from the Human Connectome Project (HCP). We further validated our observations using a separate unrelated subjects dataset from the HCP. Quantitative assessment was performed by computing tract densities and spatial overlap of identified connections between subcortical ROIs. Further, known connections between structures of the basal ganglia and thalamus were identified and visually inspected, comparing tractography reconstructed trajectories with descriptions from tract-tracing studies. Our observations demonstrate both the feasibility and reliability of using a data-driven tractography-based approach to map the subcortical connectome *in vivo*.

## 1. Introduction

A brain network is comprised of bundles of axons, which form the structural pathways (also referred to as tracts or connections), that allow transfer of information between the different regions (Sotiropoulos and Zalesky, 2019) and facilitate the performance of complex functions (Klingberg et al., 1999; Mesulam, 1998). Axons can be computationally reconstructed (represented as a streamline) using diffusion magnetic resonance imaging (dMRI), a non-invasive technique sensitive to the direction of water motion (Bammer, 2003; Conturo et al., 1999). As axons are bundled together, water molecules will preferentially diffuse parallel to the axonal trajectory, which can then be detected using dMRI to enable an *in vivo* estimation of tract trajectories. This process, known as tractography, first estimates the diffusion orientations within all imaging voxels before traversing from a starting seed location until termination criteria are met (e.g. quantitative value drops below defined thresholds) (Sotiropoulos and Zalesky, 2019). Additionally, regions of interest (ROI) can be used to define inclusion and exclusion criteria to constrain tract trajectories and facilitate identification of connections between terminal regions.

Mapping the human connectome is an important, non-trivial task that contributes to disentangling the network organization of the brain and increased understanding of changes in healthy aging or due to disease (Sporns, 2011; Toga et al., 2012). To date, much of the work studying structural connectivity using dMRI has focused on the cortico-cortical (between regions of the cortex) and cortico-subcortical (between cortex and subcortex) tracts, resulting in the development of a number of structural connectivity atlases. Such connectivity can be described as the cortical connectome. Examples of such atlases include the Johns Hopkins University white matter atlas, which identified a number of cortico-cortical white matter tracts (Hua et al., 2008; Mori et al., 2005; Wakana et al., 2007), and the Oxford thalamic connectivity atlas, which aimed to identify cortico-subcortical connectivity between regions of the thalamus and the cortex (Behrens et al., 2003a, 2003b). These atlases have been extensively used to attain an understanding of changes associated with aging as well as disease (e.g. thalamic changes in Alzheimer’s disease; Delli Pizzi et al., 2015). Validation of some of these connections have also been performed previously in studies of non-human primates (NHPs; Siwek and Pandya, 1991; Yeterian and Pandya, 1993, 1991).

Just as there are cortical connections, there is also connectivity between subcortical structures (e.g. the thalamus and basal ganglia), forming the subcortical connections, which can also be referred to as subcortico-subcortical connections. These subcortical structures are important to motor control (Gallay et al., 2008; Sommer, 2003), as well as cognition and emotion (Hollander et al., 2015). Accordingly, connections between the subcortical structures are integral and have been studied extensively in non-human primate (NHP) studies of the motor network (DeLong and Wichmann, 2007; Gallay et al., 2008; Krack et al., 2010), as well as associative and limbic networks (Alexander et al., 1986; Krack et al., 2010; Middleton and Strick, 2000; Smith et al., 1998). Previous studies have examined subcortical connections with the use of anatomical tracers, which involve injection of either anterograde or retrograde tracers at a structure of interest to map its connections. One such example involved the injection of an anatomical tracer at the ventral pallidum, which determined projections to the subthalamic nucleus (STN), as well as the hypothalamus and brainstem (Haber et al., 1993),

Studies that attempt to more comprehensively identify the subcortical connections non-invasively via tractography, that is to map the *subcortical connectome*, have been limited. The scarcity of subcortical connectome studies is in part due to the difficulty of tracking the connections in a compact region where the underlying diffusion signal is complicated by multiple diffusion orientations arising from numerous intersecting connections and structures with low anisotropy. One previous study demonstrated the ability to map connections between the basal ganglia and thalamus *in vivo* using manual segmentations before leveraging connectivity strength to parcellate the basal ganglia and thalamus into subregions (Lenglet et al., 2012). Recently, *in vivo* studies have primarily focused on individual connections that comprise specific subcortical connections and have been identified as putative targets for surgical neuromodulation (Avecillas-Chasin and Honey, 2020; Rozanski et al., 2017). In one study, the pallidothalamic tract was delineated in order to study its role in the treatment of dystonia with deep brain stimulation (DBS) (Rozanski et al., 2017), while another study examined the importance of pallidoputaminal connectivity to predict DBS outcomes also for dystonia (Raghu et al., 2021). With the aid of a number of atlas-based inclusion and exclusion ROIs, as well as extensive manual refinement, tractography has been used to identify the nigrofugal and pallidofugal subcortical connections. (Avecillas-Chasin and Honey, 2020). Recently, an attempt was made to map subthalamic tracts using *ex vivo* data, using ROIs to guide and identify specific subcortical connections (Oishi et al., 2020). All of these studies employed tractography to identify the trajectory of the connections using non-invasive techniques, highlighting a potential for tractography-guided treatment. Reliable and accurate identification of these connections has the potential to improve diagnosis and treatment options.

With reliability studies having been previously performed in tractography studies of cortical connectivity (including, but not limited to Buchanan et al., 2014; Cousineau et al., 2017; Guevara et al., 2017; Schilling et al., 2021), an evaluation of the reliability of the subcortical connectome is also warranted. Despite examination of individual subcortical connections, to our knowledge, there has yet to be a study assessing the reliability of the subcortical connectome.

Briefly, reliability is defined as the agreement of the results (e.g. similar connectivity) when applying the same methodology to different acquisitions of the same subject or to data acquired from different subjects. Not to be confused with reproducibility, another term that often gets used interchangeably, which is defined as the ability to produce similar results when using an entirely different methodology. Both are important and can provide valuable insight regarding a method or result. Reliability studies can evaluate and increase the confidence of methodological approaches used to study structural connectivity, while reproducibility studies can validate findings by comparing results produced with other techniques. In this work, we recapitulate pathways of the subcortical connectome in the Human Connectome Project (HCP) test-retest dataset. We aimed to assess the feasibility and reliability of mapping the subcortical connectome, with a specific goal of recapitulating known connections, through application of subcortical structure segmentations and probabilistic tractography. Furthermore, we sought to develop a framework that enabled evaluation of reliability for the subcortical connectome moving forward. Additional validation was performed using the unrelated subjects dataset of the HCP.

## 2. Materials and methods

Processing of the data was performed in containerized computing environments on a high performance compute cluster. An overview of the general workflow is shown in Fig. 1. Briefly, publicly available minimally pre-processed test-retest data from the Human Connectome Project was used to assess reliability of connections (identified via tractography) between subcortical structures and feasibility of identifying connections of known subcortical circuits. Analysis included evaluating tract overlap, changes in tract density, and examining identified connections with trajectories previously described in the literature. Furthermore, processing and analysis was replicated on an unrelated subset from the Human Connectome Project.

**Figure 1.**
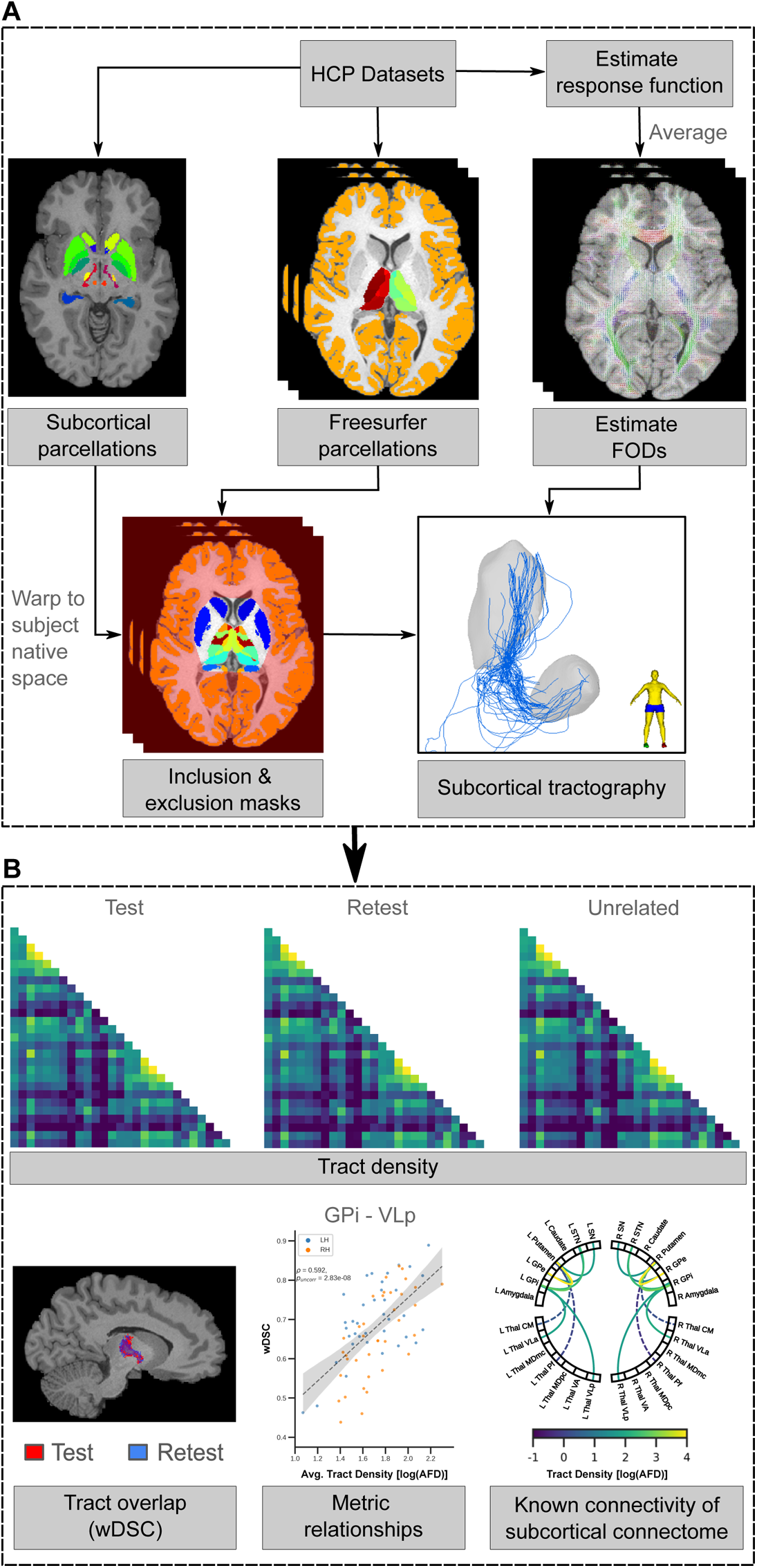
General subcortical tractography processing workflow using the minimally preprocessed HCP datasets. (A) An average response function was created from individual response functions from each acquisition (per dataset) and to estimate the FODs in each MRI session. Additionally, FreeSurfer was employed to parcellate the thalamus and obtain a cortical ribbon. Inclusion and exclusion masks were created, combining subcortical parcellations (transformed to the subject’s native space) with FreeSurfer parcellations to perform tractography on the subcortical connectome. (B) Examples of assessments performed, comparing test vs retest sessions, as well as the use of an additional unrelated dataset for further comparison.

### 2.1 Dataset

Minimally pre-processed subjects as part of the test-retest dataset (n=36; 11M/25F, aged 22-35) of the Human Connectome Project (HCP) (Glasser et al., 2013; Van Essen et al., 2013) were used to assess the reliability of subcortical connections identified via tractography. Briefly, T1-weighted (T1w) MRI scans were acquired with a 3D MPRAGE sequence (Mugler and Brookeman, 1990): resolution = 0.7 mm isotropic voxels; repetition time/echo time (TR/TE) = 2400 / 2.14 ms, while dMRI scans were acquired in opposite anterior-posterior phase-encoding directions with a pulsed gradient spin-echo sequence (Stejskal and Tanner, 1965): resolution = 1.25 mm isotropic voxels; TR/TE = 5520 / 89.50 ms; b-values = 1000, 2000, 3000 s/mm^2^ (90 directions per shell) with 18 b-value = 0 s/mm^2^ images. All data was acquired on customized Siemens Skyra 3T MRI systems (Sotiropoulos et al., 2013; Van Essen et al., 2012). Full acquisition details are described in the HCP1200 reference manual^1^. As part of the minimal pre-processing pipeline data release, all subjects underwent FreeSurfer processing (v5.3.0-HCP; Fischl et al., 2004), where the cortical ribbons were retained for further processing.

Further, subjects part of the HCP unrelated dataset that did not overlap with test-retest dataset were selected (n=85; 35M/50F; aged 22-35) for validation. Acquisition and minimal pre-processing steps from the HCP release of the unrelated dataset were identical to the test-retest dataset.

### 2.2 Regions of interest

To evaluate connections of interest, structural segmentations were used as ROIs to assist tractography generation. As previously mentioned, cortical reconstruction from FreeSurfer (Fischl et al., 2004) was first performed, retaining the cortical ribbon as an exclusion mask. In addition, subcortical structures where connections terminated were identified. Subnuclei of the thalamus were segmented using FreeSurfer (v7.1.0; Iglesias et al., 2018), while other subcortical structures (excluding the hippocampus, which is considered part of the archicortex; DeKraker et al., 2020; Duvernoy, 2013) were identified from the BigBrain subcortical atlas (Xiao et al., 2019) first registered to the MNI2009bAsym template (Fonov et al., 2011). Volumes of all subcortical structures were computed for each subject. A second exclusion mask was created from an inverted convex hull surrounding the subcortical structures to discard streamlines outside of the convex hull. FreeSurfer processing was performed in the subject’s native space, while the atlas was transformed to the subject’s native space using the Advanced Normalization Tools (ANTS v2.1.0; Avants et al., 2009). Briefly, the atlas was transformed to the subject’s native space in a 3-step process: (1) linear affine transformation, (2) non-linear symmetric normalization (SyN), and (3) HCP provided subject-specific transformations. The first two steps transform the labels from MNI2009bAsym space to MNI152NLin6Asym space, while the final step transforms the labels to the subject’s native space. Transformations between the two spaces can be found in the available repository (see data availability).

### 2.3 Tractography

All tractography processing was performed using the MRtrix3 software suite (v3.0_RC3; Tournier et al., 2019). First, individual tissue-specific response functions were estimated for each subject in both test and retest sessions using an unsupervised approach (Dhollander et al., 2016). From here, an averaged group response function was computed from the individual response functions. Fiber orientation distribution (FOD) maps were estimated for each subject with a multi-shell, multi-tissue constrained spherical deconvolution (MSMT-CSD) algorithm (Jeurissen et al., 2014), with group average response functions independently computed for the test-retest and unrelated datasets. The use of a group average response function minimizes biases in FOD maps (Raffelt et al., 2012), improving the comparability of tractography within datasets with observed differences attributed to the underlying diffusion data of an individual. Prior to performing tractography, multi-tissue informed log-domain transformed normalization was performed (Raffelt et al., 2017) on the FOD maps.

As the primary diffusion orientation is also reflected in FODs, major white matter connections (e.g. the corticospinal tract (CST)) passing through the subcortical region will hinder the ability to identify subcortical trajectories. To traverse trajectories along non-primary diffusion orientations, the iFOD2 probabilistic algorithm (Tournier et al., 2010) with a step-size of 0.35mm and maximum angle of 45° between successive steps was used. Random seeding was performed throughout the brain until 20 million streamlines, constrained to the subcortical region with the previously created exclusion mask, were selected. The chosen parameters are comparable to what is typically used in iFOD2 algorithms to perform whole-brain tractography with a noted decrease in step-size (from 0.5 × 𝑣𝑜𝑥𝑒𝑙 𝑠𝑖𝑧𝑒 to 0.25 × 𝑣𝑜𝑥𝑒𝑙 𝑠𝑖𝑧𝑒) to sample more frequently along a streamline’s trajectory.

Following tractogram creation, each streamline was assigned a weighting to reflect its contribution to the underlying diffusion signal using the updated spherical-deconvolution informed filtering of tractograms (SIFT2) technique enabling the assessment of tract densities (Smith et al., 2015). Using MRtrix3, structural connectivity was established by identifying the nearest subcortical label within a 1.5mm radius at each terminal end of a given streamline. Due to the low anisotropy within gray matter, streamlines whose trajectories intersect other subcortical labels prior to reaching the terminal structures were discarded (see Discussion). Furthermore, the CST, which represents a dominant tract passing in proximity to many subcortical connections, was separately identified in order to visually assess its influence on derived tracts. Identification of the CST was performed using the brainstem and segmentations of both pre- and post-central gyri identified by FreeSurfer as inclusion regions of interest. Generation of the CST was performed until 500 streamlines were identified in each hemisphere. Similar to the connectivity of the subcortical connectome, streamlines had to terminate within a 1.5mm radius of these segmentations to be considered a part of the CST.

### 2.4 Assessment of reliability and accuracy

An investigation into known subcortical connections of the motor, limbic, and associative networks was performed, quantitatively assessing reliability of tract densities and spatial overlaps of identified connectivity. Connectivity between structures associated with the networks were identified and extracted (DeLong and Wichmann, 2007; Gallay et al., 2008; Krack et al., 2010), with both ipsilateral self-connections (i.e. tracts that start and end in the same ROI) and *inter-* hemispheric connections excluded from analysis. Further, subcortical connections that connect to thalamic nuclei on both terminal ends were also excluded. Visual inspection of known connections of the subcortical connectome was also performed to evaluate accuracy of tractography-produced trajectories with previously described literature.

#### 2.4.1 Anatomical assessment

Using the method employed to identify connectivity between subcortical structures, a large number of potential connections were found. Since our goal was to recapitulate known subcortical connections with *in vivo* tractography, we focused on those that have been well described in the literature depicting motor, associative, and limbic subcortical circuitry (DeLong and Wichmann, 2007; Gallay et al., 2008; Krack et al., 2010). Connectivity between subcortical structures of the basal ganglia and thalamus were both visually and quantitatively examined, evaluating tract trajectories, densities, and overlap.

#### 2.4.2 Tract density

Streamlines weighted by their contribution to the underlying diffusion signal were summed to calculate the tract density (also referred to as apparent fibre density (AFD); Raffelt et al., 2012) of the connection between two subcortical structures. A connectivity matrix for each subject was created with the AFD representing the edge strength between two ROIs (nodes). Further, the percent change in AFDs were calculated between test and retest sessions using equation 1:

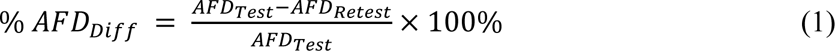

Additionally, intraclass correlation (ICC) was computed for the tract densities between the two datasets as a metric of consistency using a two-way, mixed effects model (McGraw and Wong, 1996). Prior to computing an ICC, an analysis of covariance (ANCOVA) was first performed to identify and account for covariates (age, subject motion, brain volume) with a significant effect on the tract density via linear regression. In this model, the “raters” (column factor) were the corrected tract densities and the “targets” (row factor) were the test and retest session connectivity. A paired t-test was also conducted between average AFDs of the test and retest sessions. To compare average connectivity of the basal ganglia with average connectivity between the basal ganglia and thalamus, an one-way ANOVA was performed. The impact of ROI volume on AFD was assessed using ordinary least squares multiple regression, treating the average ROI volume across subjects as an independent variable and AFD as the dependent variable. Further, Spearman’s correlation was performed between average AFD and the absolute percent change between test and retest sessions.

#### 2.4.3 Voxel-wise spatial overlap

An AFD map was first created for each tract identified in the test and retest sessions by identifying streamlines passing through each voxel. The sum of streamline weights were assigned to corresponding voxels. Following assignment of streamline weights, the fraction of the tract (a value between 0 and 1) passing through a voxel is determined from the AFD map and used to compute the overlap between tracts from the weighted Dice similarity coefficient (wDSC; Cousineau et al., 2017). Briefly, the wDSC is a modified Dice similarity coefficient for assessing tractography overlap, minimizing the penalization applied to streamlines further from the core of the tract (Cousineau et al., 2017). The wDSC is computed from equation 2:

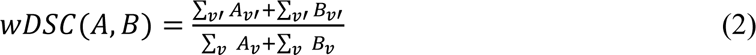

where A and B represent the fraction of streamlines (between 0 and 1) passing through a voxel and v’ represents a corresponding non-zero voxel in A and B. The numerator of equation 2 computes the sum of overlapping non-zero voxels between A and B, while the denominator calculates the total sum of non-zero voxels in A and B respectively. Computed overlaps from wDSC follow similar indicators of agreement as the conventional Dice similarity coefficient: poor (<0.2), fair (0.2 – 0.4), moderate (0.4 – 0.6), good (0.6 – 0.8) and excellent (>0.8; Kreilkamp et al., 2019).

In addition to comparing the tract overlap, a Spearman’s correlation was also computed between the average AFD across the two datasets. Similar to AFD, a one-way ANOVA was performed to compare the wDSC for connectivity of the basal ganglia with connectivity between the basal ganglia and thalamus.

#### 2.4.4 Identifying a connectivity threshold

The previously computed connectivity matrices were thresholded in an effort to retain the most reliable tracts, while discarding potential false positive connectivity. As noted in previous studies, defining a threshold is a non-trivial task (Shadi et al., 2016; Zhang et al., 2018). If the chosen threshold is too low, tracts that are not reliably identified may remain, including those that do not exist in reality, but if it is too high, legitimate connections may be discarded. Common approaches include choosing an arbitrary threshold such that the majority of the subjects to be analyzed retain the same connections (Li et al., 2009) or by sweeping through a range of thresholds (Li et al., 2012). More recently, a test-retest metric was proposed, wherein reliability was evaluated across a range of thresholds and a final threshold was selected where the change was at a minimum (Zhang et al., 2018). Here, we selected our threshold by following a similar test-retest reliability procedure, using the tract overlap (wDSC) as the reliability measure. We first stepped through a range of AFD values to threshold the connectivity matrix before calculating the wDSC for each thresholded matrix. Additionally, we computed the change in average wDSC between each step across the range of AFD values. The wDSC threshold is selected at the first occurrence where the change between steps is 0 and identified the corresponding average AFD threshold. Supplementary Fig. 1 demonstrates examples of the connectivity matrix at different thresholds of tract overlap.

#### 2.4.5 Validation with unrelated dataset

Processing and analysis of the HCP unrelated dataset followed the same workflow as before with the test-retest dataset. As before, known connectivity of the subcortical connectome was both visually and quantitatively assessed. AFD matrices were computed for each subject as before, and further separated by hemispheric connectivity to compare with previous findings. With only a single acquisition session in the unrelated dataset, an average AFD matrix was computed across subjects, and a Pearson’s correlation was performed against the average AFD matrices of the test and retest sessions to evaluate the similarity of the subcortical connectome. As with the test-retest dataset, the relationship between AFD and the size of the subcortical structures was also evaluated.

## 3. Results

### 3.1 Networks of the subcortical connectome

We investigated the ability of *in vivo* tractography to both identify and reliably reproduce the connectivity between different acquisitions of the same human subject, focusing on known subcortical connections of the motor, associative, and limbic circuits (DeLong and Wichmann, 2007; Gallay et al., 2008; Krack et al., 2010).

#### 3.1.1 Identification of known subcortical connections

Motor network connectivity using the described tractography methods could successfully recapitulate known connections as previously described in the literature (Fig. 2). Similarly, known connections of both the associative and limbic network connectivity were also successfully captured (Supplementary Fig. 2A and Supplementary Fig. 2B respectively). A wDSC of 0.58 was selected as the final overlap threshold, which corresponded to a AFD threshold of 6.5 AFD. Of the known connectivity comprising the motor network, 78% (14 out of 18) of the identified connections met the threshold. In the associative and limbic networks, 100% and 79% (11 out of 14) of the observed connectivity met the AFD threshold respectively. Connectivity failing to meet this threshold was commonly found between a thalamic nucleus (which was often small) and another subcortical structure (see Supplementary Table 1 for full details), for example, between the putamen and the centromedian and parafascicular nuclei of the thalamus in both test and retest sessions. Connectivity between the globus pallidus internus (GPi) and either division of the mediodorsal nucleus of the thalamus failed to meet the AFD threshold in both test and retest sessions (both hemispheres for the magnocellular division and left hemisphere for the parvocellular division).

**Figure 2.**
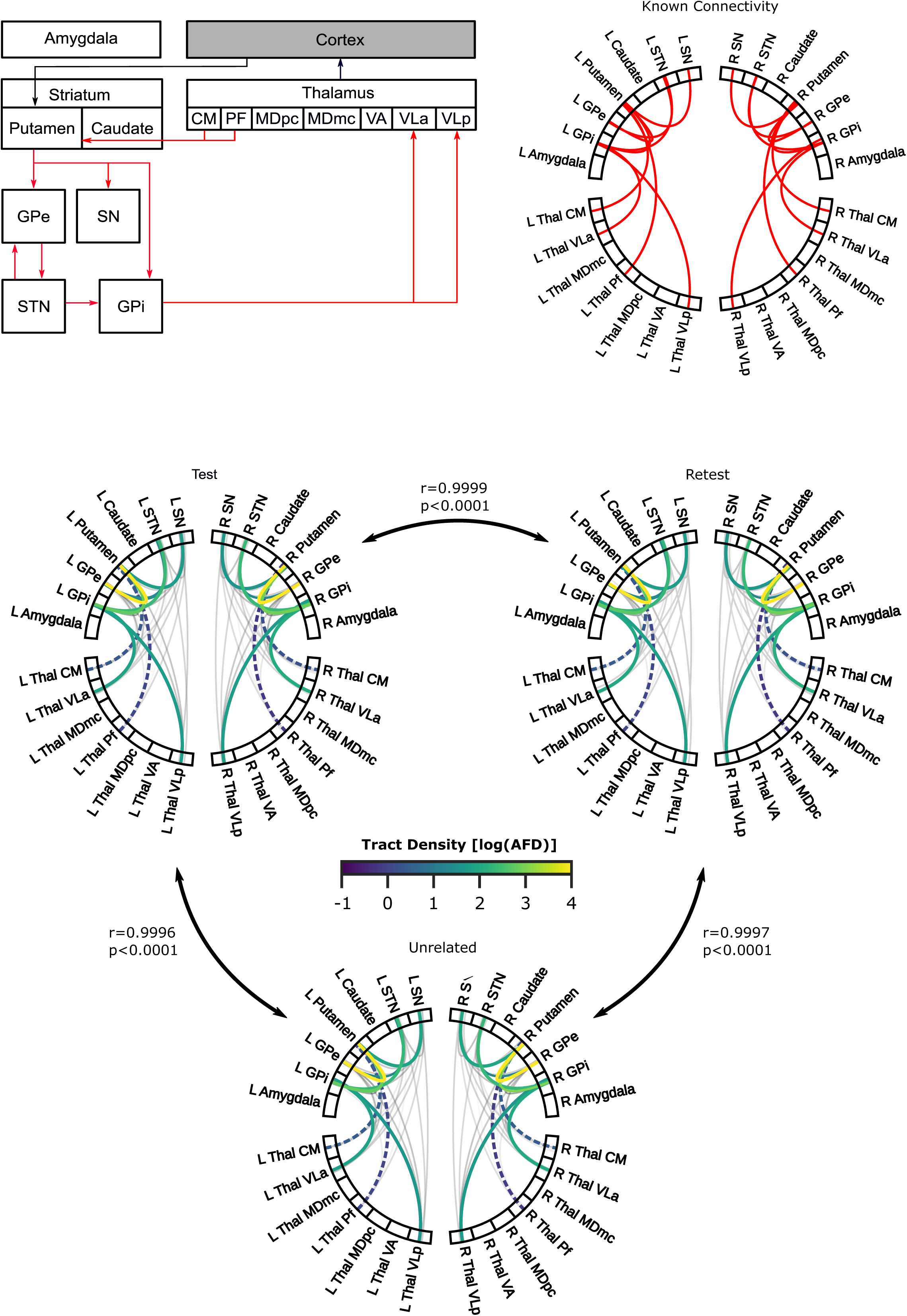
Diagram of known anatomical subcortico-subcortical connections (in red) of the motor network. (A) Connections identified from literature are depicted in a diagram (left) and chord plot (right). (B) Chord plots exhibiting average log-transformed tract densities from tractography derived connections are displayed for test-retest (top-left, top-right) and unrelated (bottom) datasets from the Human Connectome Project. Coloured lines represent known connections, with dashed coloured lines specifically indicating known connections that did not meet the selected tract density threshold. Grey lines denote connections identified from tractography, but not identified in tract-tracing literature. Pearson correlations between datasets are shown next to the comparison indicators.

Identified connections were also visually inspected, examining the connected structures and their trajectories. In observations of tract density, it was previously noted that basal ganglia connections (e.g. non-thalamic ROI to non-thalamic ROI) were denser, while connections between the basal ganglia and thalamus (e.g. non-thalamic ROI to thalamic ROI) were sparser. Visual inspection of the known trajectories, reflected the previous observation of denser connectivity between basal ganglia structures, which are also shorter and more direct. Conversely, connections between the basal ganglia and thalamus were sparser with longer and more curved trajectories. These longer trajectories increased the potential for intersecting GM structures between the basal ganglia and thalamus as was the case for connections between the ventrolateral anterior nucleus of the thalamus (VLa) and GPi (Fig. 3), as well as between STN and globus pallidus externa (GPe) / GPi (Fig. 4). It was observed that certain thalamic nuclei were more difficult to reach, as trajectories would have to pass through other surrounding thalamic nuclei. Some spurious streamlines were also noted (e.g. streamlines that looped in the brainstem). Full descriptions of known subcortical connections can be found in Supplementary Table 1.

**Figure 3.**
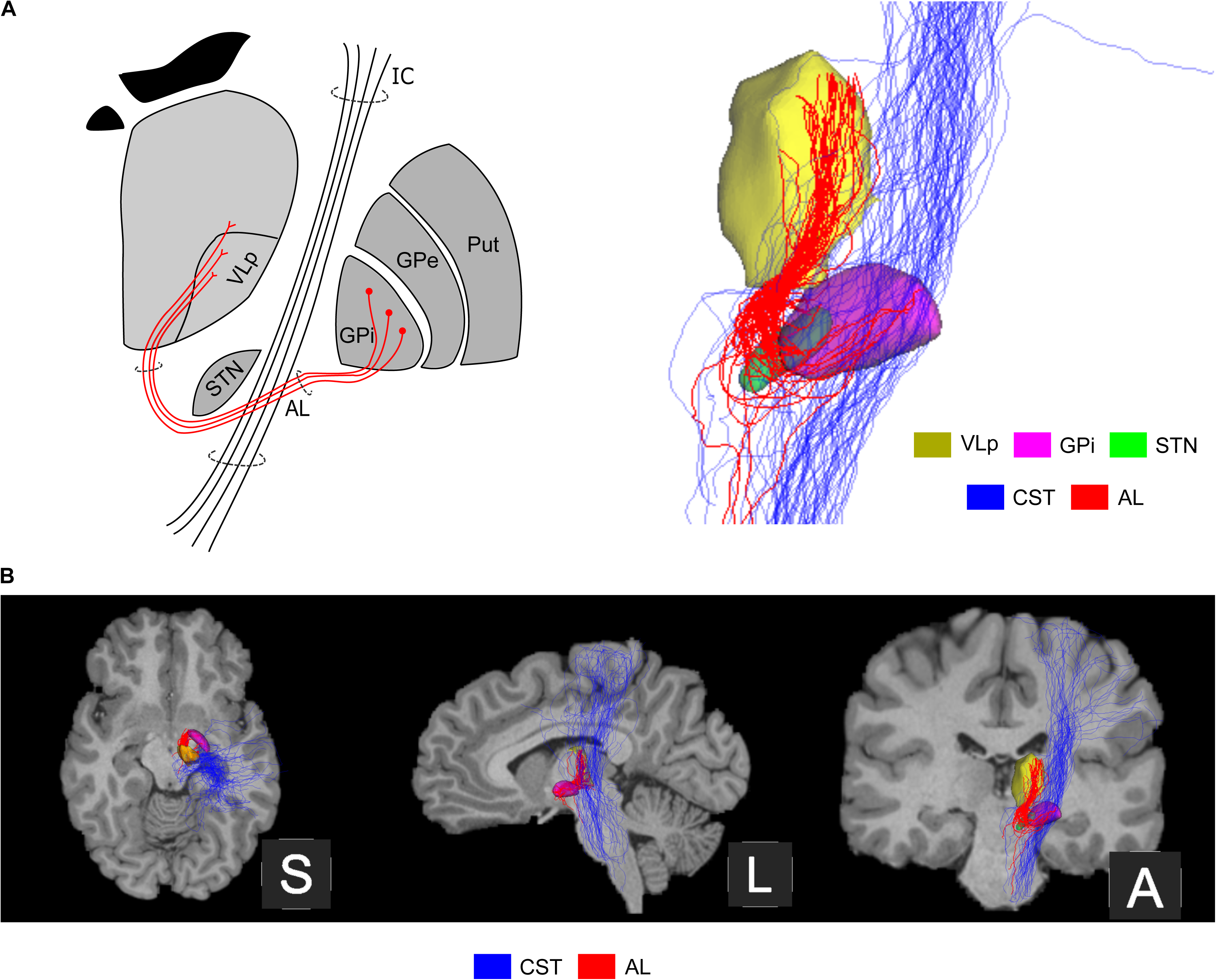
A single subject example of a connection found between the internal segment of the globus pallidus (GPi) and ventrolateral posterior nucleus of the thalamus (VLp). Manual refinement of tractography and construction of full corticospinal tract trajectory was performed for visualization purposes. (A) Depiction of ansa lenticularis (AL) from the tract tracing literature (left), compared with tractography identified trajectory (right) viewed from coronal anterior. The CST is also displayed to demonstrate the major WM tract passing through. (B) Three views (from left to right): superior, sagittal left, and coronal anterior exhibiting the trajectories of AL and corticospinal tract (CST) overlaid on a T1-weighted anatomical image.

**Figure 4.**
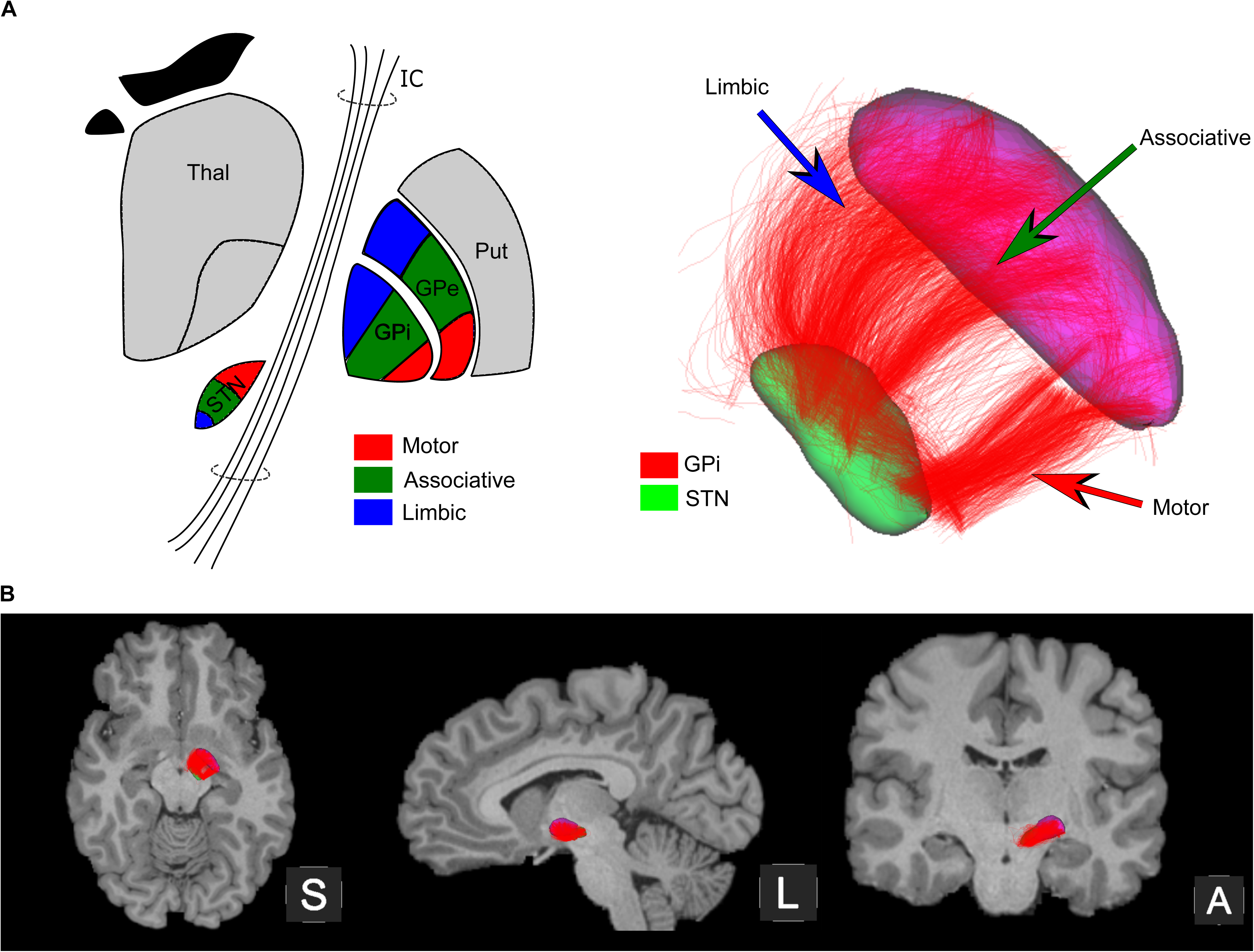
A single subject example of a connection found between the globus pallidus internal segment (GPi) and subthalamic nucleus (STN). For visualization purposes, refinement of tractography was performed and opacity was reduced to 20% to highlight the different segments observed. (A) Depiction of the approximate regions associated with different networks (e.g. motor, associative, limbic), identified from the literature (left) is shown for the STN and both internal and external segments of the globus pallidus: motor (red), associative (green), limbic (blue). Tractography identified trajectories (right) between the STN and GPi are shown from an inferior-to-superior (ventral) view, highlighting the different components associated with each network. (B) Three views (from left to right): superior, sagittal left, and coronal anterior exhibiting the connectivity between GPi and STN overlaid on a T1-weighted anatomical image.

#### 3.1.2 Reliability of known subcortical connections

The reliability of identified connections was evaluated via tract overlap within motor, associative, and limbic networks. Connections between basal ganglia structures exhibited good overlap (average wDSC = 0.751 and 0.722 for left and right hemispheres respectively), while connections between the basal ganglia and thalamus demonstrated moderate to good overlap with the VLa (average wDSC = 0.543 and 0.560 for left and right hemispheres), ventrolateral posterior nucleus of the thalamus (VLp; average wDSC = 0.527 and 0.451 for left and right hemispheres), and the ventroanterior nuclei of the thalamus (VA; average wDSC = 0.576 and 0.629 for left and right hemispheres), which all had boundaries in the easier to reach lateral region of the thalamus. Some of the connections to the thalamus in each network exhibited poor overlap (average wDSC = 0.176 and 0.167 for left and right hemispheres), coinciding with the same ones that demonstrated a low AFD. For connections between basal ganglia structures, a poor to moderate overlap was only found between the caudate and amygdala (average wDSC = 0.354 and 0.152 for left and right hemispheres), where the tract was sparse and trajectories would have had to pass through other GM structures (e.g. putamen, GPe, GPi). Additionally, lower overlap was observed in the connections between the basal ganglia and thalamus, in particular connections to the mediodorsal nuclei of the thalamus (MD), which was more difficult to reach and in which trajectories also had to potentially traverse other nuclei of the thalamus. Despite the overlap observed in a few connections, good overall reliability was demonstrated for connectivity of each network, with similar measurements for each hemisphere.

### 3.2 Evaluation of subcortical connectivity matrices

Connectivity matrices were created for both test and retest sessions between subcortical structures for all subjects. Visual assessments were first performed, followed by quantitative evaluation of all intra-hemispheric subcortical connections. Reliability of the subcortical connectome was also evaluated and noted to be similar to what was previously assessed for known connections.

#### 3.2.1 Tract density of all intra-hemispheric connections

Connectivity matrices for test and retest sessions were created from the computed AFD between subcortical structures for all subjects. A visual assessment of the computed matrices was first performed, followed by a quantitative evaluation of the computed AFDs. Matrices were observed to be similar across subjects and test-retest sessions (Supplementary Fig. 3). Connections between basal ganglia structures were often denser (AFD = 2.33 log(AFD) and 2.24 log(AFD)) in left and right hemispheres respectively, averaged across test and retest sessions) than connections between the basal ganglia and thalamus (AFD = 1.34 log(AFD) and 1.26 log(AFD)) in left and right hemispheres respectively, averaged across test and retest sessions). The difference between the two groups was also corroborated with a one-way ANOVA for both the left (F = 19.19, p < 0.05) and right (F = 3.75, p < 0.05) hemispheres. Fig. 5A demonstrates the average tract densities across the test-retest session.

**Figure 5.**
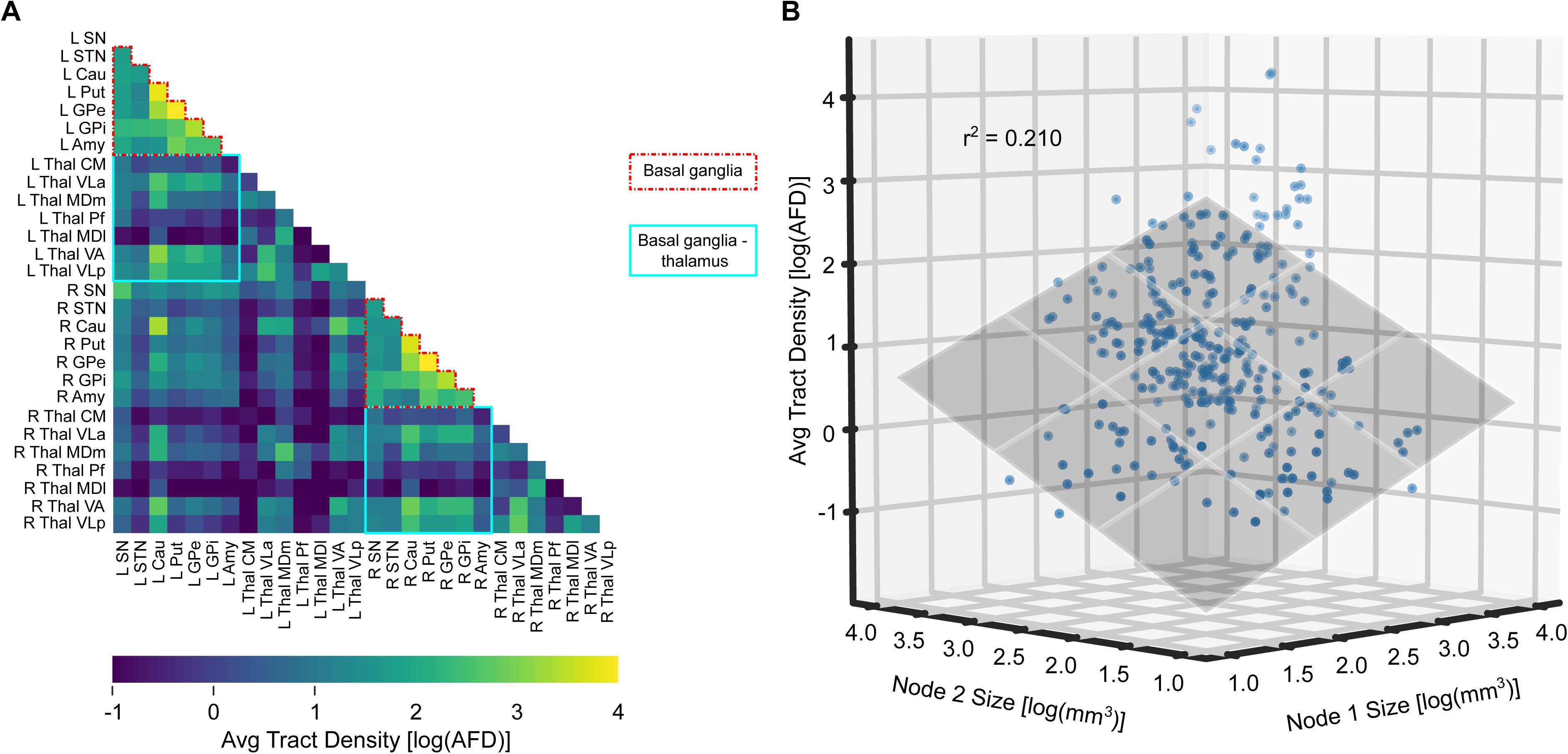
Averaged log-transformed tract densities across the test-retest dataset. (A) Connectivity matrix highlighting the two groups of connections observed: basal ganglia (red) and basal ganglia - thalamus (blue). (B) The relationship between the average tract density of connections and the volume of the terminal nodes is shown in a scatterplot. Tract density was noted to increase with an increase in volume of at least one terminal structure.

The influence that the volume of terminal subcortical structures had on AFD was also assessed. By plotting the average AFD against the volume of the two terminal subcortical structures (Fig. 5B), the average tract density was observed to increase as the volume of one of the two structures increased. Performing an ordinary least squares regression, we identified a positive linear relationship between the AFD and the size of the two subcortical structures (r^2^ = 0.210, p < 0.05).

#### 3.2.2 Reliability of all intra-hemispheric connections

Using the previously computed connectivity matrices, an average AFD matrix across subjects was created for the test and retest sessions independently (Fig. 6A). Individual subject matrices, as well as average session matrices were visually inspected, and minimal differences were observed between test and retest sessions. A linear relationship was identified between the AFDs of the test and retest sessions (Supplementary Fig. 3A; ρ = 0.997, p < 0.05), with greater variability observed between sessions when the AFD was low. Connectivity was further divided by hemisphere (i.e. *intra-*hemispheric left vs right) and a box plot was created to visually compare test and retest tract densities (Fig. 6B). No differences in hemispheric connectivity were observed between test and retest sessions, which was further corroborated after performing a paired t-test between average test and retest AFDs (t=1.52, p = 0.264 and t=1.06, p = 0.293 for left and right hemispheres respectively).

**Figure 6.**
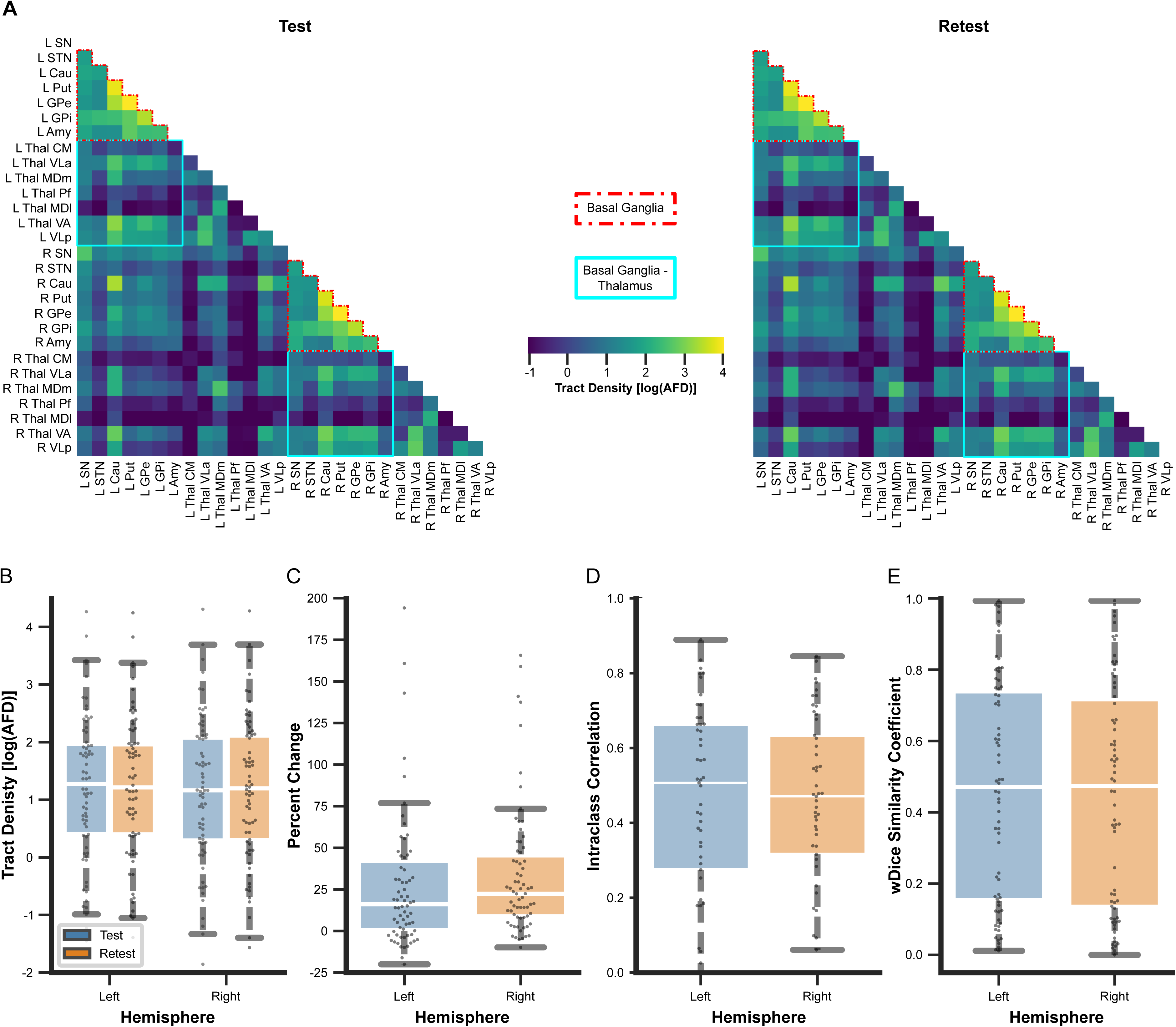
(A) Test (left) and retest (right) connectivity matrices are shown, visualizing the log-transformed tract densities between subcortical structures. (B) Log-transformed tract densities of the basal ganglia and between basal ganglia and thalamic connectivity are plotted, separated by hemisphere and session. (C) Percent change of tract densities between test and retest sessions, separated by hemisphere. (D) Intraclass correlations, measuring consistency between sessions, are shown, separated again by hemisphere. (D) wDSC, assessing spatial overlap between sessions, plotted by hemispheric connectivity. For all boxplots, the middle line marks the median metric, while whiskers define the maximum and minimum values of each metric, excluding outliers.

To quantify the consistency of AFD between test and retest sessions, we computed the percent change of corresponding subcortical connections between sessions finding on average a percent change of 36% and 32% for the left and right *intra*-hemispheric AFD respectively (Fig. 6C). As previously noted, in test-retest pairs where AFD was low, greater variability was observed. Correspondingly, a greater absolute percent change was more likely to be associated with a sparser connection. A Spearman’s correlation between the absolute percent change of AFD and the average density across test-retest sessions demonstrated a negative correlation (Supplementary Fig. 3B; ρ = -0.240, p < 0.05), indicating decreasing percent change as tract density increased. Further, an average intraclass correlation (ICC) of 0.50 and 0.48 was computed for left and right hemispheric connectivity respectively between test and retest AFDs after performing a linear regression to account for brain volume, a covariate identified to demonstrate a significant effect (F = 7.068, p < 0.05) after performing an ANCOVA (Fig. 6D). As observed in the tract density, ICC was noted to be greater in connections between basal ganglia structures (ICC = 0.58 and 0.49 for left and right hemispheres respectively) than between the basal ganglia and thalamus (ICC = 0.46 and 0.47 for left and right hemispheres respectively).

Voxel-wise spatial overlap of tracts (calculated via wDSC), was also computed between test-retest pairs as another measure of reliability. We observed an average wDSC of 0.46 and 0.45 for the left and right *intra-*hemispheric connectivity respectively. We also plotted and performed a Spearman’s correlation between wDSC and average AFD across both sessions where wDSC is expected to increase with AFD before plateauing. As expected, we identified a sigmoid relationship (Supplementary Fig. 3C; ρ = 0.950, p < 0.05) between the two (Fig. 3E), with good overlap achieved at a AFD around 2.0 log(AFD) and reaching maximum overlap at approximately around 2.5 log(AFD). The overlap remained low while the AFD was less than 0 log(AFD) before slowly increasing until the overlap began to peak at a log-transformed AFD of around 2 log(AFD). As wDSC was highly correlated with AFD, we further separated and evaluated the wDSC to the two previously identified groups. As with observations from known subcortico-subcortical connections, we noted better overlap in basal ganglia connectivity (average wDSCs = 0.75 and 0.72 for left and right hemispheres respectively) than in connectivity between the basal ganglia and thalamus (average wDSCs = 0.46 and 0.44 in left and right hemispheres respectively). The difference observed between the two groups was also supported with a one-way ANOVA for both the left (F = 33.53, p < 0.05) and right (F = 41.64, p < 0.05) hemispheres.

### 3.3 Observations in HCP unrelated dataset

An identical analysis was performed on a subset of the HCP unrelated dataset, where similar observations were noted. An average connectivity matrix was computed and compared against the test-retest dataset, where notably a Pearson’s correlation coefficient of 0.99 was demonstrated against both the test (p < 0.05) and retest (p < 0.05) sessions, indicative of highly similar AFDs with the unrelated dataset. Full details of the results from this validation can be found in the Supplementary Material.

## 4. Discussion

In this study, we evaluated both the feasibility and reliability of identifying the subcortical connectome using *in vivo* tractography data, specifically evaluating the possibility of recapitulating known connections from classic studies of the subcortex. We demonstrated the ability to identify most of the subcortico-subcortical connections (39 out of 46, 85%) described in the literature. Furthermore, we were able to demonstrate test-retest reliability and replicate this analysis on a separate HCP subset, observing near identical results (see Supplementary Data) and again recapitulating most known connections (38 out of 46, 83%). In the following subsections, we compared our observations with the existing literature. Importantly, we also recount the challenges that were faced in studying the subcortical connectome *in vivo* and suggest possible solutions.

### 4.1 Identification of known subcortical connections

Tract-tracing studies have been performed in NHPs to identify and study networks of the subcortical connectome (Parent and Carpenter, 1996; Sato et al., 2000a, 2000b; DeLong and Wichmann, 2007; Gallay et al., 2008; Krack et al., 2010; Haber et al., 1993). A few of these tracts that comprise the networks have also been identified as potentially important neuromodulatory targets (Rozanski et al., 2017; Avecillas-Chasin and Honey, 2020; Raghu et al., 2021; Haber et al., 2021; Tsuboi et al., 2021) and may also be critical biomarkers in aging or disease progression (Abos et al., 2019). In the present study, we identified and investigated connections of the motor, associative, and limbic networks observing that most known subcortical connections could be recapitulated with a data-driven probabilistic tractography approach. The identified connections were visually inspected to evaluate the trajectories and their plausibility, comparing our observations with the literature. The ease of identifying trajectories between structures varied, with proximity between terminal structures and density playing a factor. Some notable observed trajectories included connections between the GPi and both the VLa and VLp nuclei of the thalamus, as well as between the STN and both GPe and GPi (Fig. 3 and Fig. 4), which have both been well studied. Full details regarding our observations are found in Supplementary Table 1.

First, focusing on the connection between GPi and VLa/VLp, one plausible trajectory observed was the ansa lenticularis (Fig. 3). Similar origins of the ansa lenticularis observed in this study from tractography reconstruction have been previously described from tract-tracing studies in NHPs, with similar projections from the GPi to the VLa and VLp (Gallay et al., 2008). The connection has also been described by (Parent and Carpenter, 1996), noting a trajectory that “forms a well-defined bundle on the ventral surface of the pallidum…” curving around the posterior limb of the internal capsule before continuing posteriorly. Along this trajectory, the ansa lenticularis is known to converge with the lenticular fasciculus in the fields of Forel to form the thalamic fasciculus, which continues to VLa and VLp. While we were able to note the termination in the VLa and VLp in our observations, we were unable to delineate the transition from ansa lenticularis to thalamic fasciculus. Further, sparse connections were observed to cross the region of the internal capsule to connect the GPi with VLa and VLp, which may be part of the lenticular fasciculus.

Another notable connection observed was between the STN and globus pallidus, including both the internus (GPi) and externus (GPe) segments (Fig. 4). Direct trajectories were seen, with noticeable separation differentiating trajectories between the motor, associative, and limbic regions of each structure. Similar separations were also observed in a connectivity-based segmentation by (Bertino et al., 2020), who noted an anteroposterior axis arrangement of the limbic, associative, and sensorimotor regions to the GPi and GPe. We observed sparse connections to the associative region of the GPe, attributed to a combination of the presence of the GPi and the constraints imposed on tracts going through wayward GM structures. Nonetheless, similar termination was not only observed in the previously mentioned study (Bertino et al., 2020), but also in tract-tracing studies, with limbic areas of the STN forming connections with the limbic region of the pallidum and likewise for associative and sensorimotor connections (Karachi et al., 2005). Other tract-tracing studies have noted similar connections (Parent and Carpenter, 1996; Sato et al., 2000b, 2000a), suggesting projections from STN to GPe, such as those observed from tractography.

Constraints were imposed to minimize the presence of false positive connections including the use of a convex hull, exclusion of wayward GM structures, and applying an AFD threshold. While these constraints did not completely eliminate false positive connections, only a small number were observed relative to the total AFD between subcortical structures (see section 3.1 and Supplementary Table 1). In the context of evaluating trajectories, false positives were identified as streamlines with implausible trajectories (e.g. crossing the mid-sagittal plane or coursing into the CSF) or those part of major white matter connections (e.g. CST). We noted spurious tracts of the CST, the dominant wayward tract traversing the subcortex (Johnson et al., 2008), which was falsely included as streamlines in a number of connections between subcortical structures. The presence of the CST is a result of the dominant diffusion signal, complicating the ability to accurately identify subcortical connections. Consequently, some streamlines predominantly follow the orientation of the dominant diffusion signal until reaching the boundary of the convex hull, where they continue by following its boundary due to the imposed exclusion criteria. Spurious streamlines were also observed to form a loop projecting back towards the cortex after entering the brainstem, where connectivity is expected to traverse from subcortical structures (Haber et al., 1993). This is likely caused by a combination of the convex hull exclusion mask and the lack of meeting a termination criteria as the streamline traverses down towards the brainstem. Careful inclusion of additional constraints, such as segmentations from *a priori* anatomical knowledge, may be useful to aid in the removal of these spurious streamlines. Truncation of a streamline once it reaches the boundary is one possible solution instead of waiting for a termination criteria to be met.

### 4.2 Reliability of the subcortical connectome

Upon visual inspection, the connectivity matrices demonstrated subjectively similar tract densities (AFDs), that is the sum of weighted streamlines that comprise a connection, across subjects and datasets. For a given subject in the test-retest dataset, the tract density was expected and observed to be similar across sessions. After correcting for brain volume, a covariate with significant effect on tract density, this was reinforced quantitatively, where no significant difference was identified between test-retest subjects and an average ICC of 0.49 and 0.48 was observed for left and right hemispheric connectivity respectively, with a noted higher reliability in connections between basal ganglia structures (see section 3.1.2). While AFDs were not perfectly identical between sessions, changes in AFD were likely due to differences in the acquired data between sessions. Furthermore, tractography seeding was performed randomly within the brain mask until the desired number of streamlines were met. The SIFT technique employed in the present study helped to minimize the differences between observations by weighting each streamline to best match the underlying diffusion signal (Smith et al., 2015). Comparison of AFD reliability with other studies was difficult as different metrics of density are often employed (e.g. raw streamline count) and further compounded by the limited number of subcortical connectome studies (Lenglet et al., 2012). In a comparison with a study of the cortical connectome, AFDs weighted by length were employed to examine the consistency of connections of interest (Buchanan et al., 2014). In their assessment, a similar average ICC of 0.62 was demonstrated, suggesting our findings within the subcortical connectome are comparable.

To validate our findings, we also replicated the analysis on the HCP unrelated dataset, observing similarities between the two datasets from visual inspection and quantitative comparison. We demonstrated high similarity of connectivity matrices across datasets, with a Pearson’s correlation coefficient of 0.99 between the averaged connectivity matrix of the unrelated dataset as well as test and retest connectivity matrices. Due to a lack of subcortical connectome reliability studies, direct comparison of our findings was challenging as often different connectomes were investigated, typically focused on cortico-cortical or cortico-subcortical connections. However, in an investigation of cortical connectome reliability across different different resolutions, Pearson’s correlation coefficients between 0.724 (high resolution) to 0.958 (low resolution) were computed between connectivity matrices of different subjects (Cammoun et al., 2012). While we acknowledge there were differences in the acquisition and protocol, our observations suggest that the subcortical connectome can be reliably reconstructed to a similar degree as the cortical connectome.

In addition to being able to reliably reconstruct similarly dense connections, it is also important to be able to capture the trajectory of the connections in a reliable manner. To that end, we computed the wDSC to measure the voxel-wise overlap of identified connectivity between test and retest sessions, minimizing the penalty on streamlines further from the tract core. Connections with similar trajectories would traverse the same voxel space and consequently demonstrate higher wDSCs. To our knowledge, while no previous work has evaluated tract overlap of the subcortical connectome, wDSC has been used to demonstrate the reproducibility in the cortical connectome (Boukadi et al., 2019; Cousineau et al., 2017). Our observed wDSC in connectivity between basal ganglia structures was within the reported range (wDSC = 0.71 to 0.82) of four examined cortico-cortical tracts identified using similar techniques (Boukadi et al., 2019). Similarly, wDSC has been employed to examine test-retest reliability of cortico-cortical tractography in the Parkinson’s Progression Markers Initiative dataset, where a wDSC of 0.72 was identified as a threshold for good overlap in their study (Cousineau et al., 2017). The same study also examined reliability of cortico-subcortical connections using an ROI defining a general cortical region (e.g. sensorimotor cortex, associative cortex, limbic cortex) to an ROI defining a subcortical structure (e.g. caudate, putamen, thalamus), where poor reliability of cortico-subcortical connectivity was noted. The poor reliability was attributed to a combination of the quality of atlas used to define ROIs, partial volume effects, motion and the low resolution of the data, as well as the difficulty of performing tractography to the subcortical brain regions where structures are generally smaller (Cousineau et al., 2017).

Overall, we demonstrated that the methods employed in the present study can produce subcortical connectomes with comparable reliability to those that have been used to study cortico-cortical and cortico-subcortical connectomes. Although we observed worse reliability in connectivity between the basal ganglia and thalamus, we believe this is primarily attributed to the sparse connections resulting from imposed GM constraints. Further some nuclei of the thalamus were more difficult to reach, with other nuclei present along the expected trajectory, while others were smaller in size. Some changes to the tractography algorithm (e.g. angle, maximum streamline length, etc.) or further optimization of constraints may be required to improve the overall connectivity with the thalamus. In subsequent work, the described framework can be leveraged to evaluate the impact of modifications to the original algorithm and their impact on the resultant tractography.

### 4.3 Clinical significance

The ability to reliably identify subcortical connections has important clinical implications for diagnosis and treatment planning, potentially improving targeting of specific subcortical structures. Previous studies have examined neuromodulation of specific subcortical connections (Avecillas-Chasin and Honey, 2020; Rozanski et al., 2017). Accurate and robust identification of the subcortical connectome can facilitate and enhance the ability to study pathologic changes due to disease. Furthermore, the ability to reliably identify subcortical connections also increases the likelihood of avoiding collateral connections, which can result in undesirable side effects. One consideration for clinical translation is the acquisition protocol and scan time. While clinical data is typically collected at lower angular and spatial resolutions than that of the data in this study, recent advancements in parallel imaging will help to make higher quality diffusion MRI feasible in a clinically-feasible time frame.

### 4.4 Implementation choices for the tractography algorithm

Tractography involves choices that need to be made at each step of the workflow that affect downstream steps and analysis. One such decision was the choice of segmentations used to identify connectivity of the subcortical connectome. As we were interested in the connectivity between specific subcortical structures, we pooled together existing atlas-based segmentations (Iglesias et al., 2018; Lau et al., 2020; Xiao et al., 2019) to serve as terminal ROIs and to minimize variability that may be introduced by manual segmentation. Choice of atlas-based segmentation was influenced by convenience and familiarity, with a focus on tools that are openly available. The thalamus labels used are readily available through a commonly used neuroimaging software package (i.e. FreeSurfer; Iglesias et al., 2018). Openly available segmentations of other subcortical structures were propagated from a single high-resolution template (Xiao et al., 2019). Different approaches to identifying ROIs can lead to varying shapes and boundaries, which would have some downstream effects on identified connectivity. For example, in histology-based segmentations, ROIs may be defined by the underlying nuclei, whereas in structural connectivity-based segmentations, ROIs may be related to regional connectivity. Furthermore, certain structures, including the caudate or putamen can also be subdivided into different components (Khan et al., 2019; Tian et al., 2020; Tziortzi et al., 2014), similar to the thalamus. With many segmentation schemes readily available and multiple considerations to contemplate, it is important to note that choice of segmentation is often dependent on the aims of the specific study (Arslan et al., 2018).

Segmentation accuracy is also important for capturing the true underlying subcortical structure, with size influencing the reconstructed connectivity as has been previously noted (Sotiropoulos and Zalesky, 2019) and also observed in this study. An ROI larger than the structure, especially in the small subcortical region, may overlap with other structures or extend into the WM or CSF. On the other hand, an ROI smaller than the structure may exclude connectivity that does not reach the boundaries, although some of this is alleviated by using the radial search strategy employed. Due to the relationship between ROI size and tract density (see Fig. 5B), the ability to identify connections in small structures is challenging and some expected connections may remain unidentified. Nonetheless, the segmentations present incorporated data from histology and largely reflect the underlying anatomy.

We chose to include a GM exclusion criteria in our tractography algorithm, removing connectivity passing through other subcortical structures along its trajectory. This choice was made in part to limit the number of false positives passing through GM structures where multiple diffusion orientations and low anisotropy are often observed that result in a significant increase in spurious streamlines. However, it is known that subcortical connections can pass through other structures (Sato et al., 2000a, 2000b). As a result of this constraint, we noticed sparse connectivity between regions where another GM structure is along the expected trajectory. One possible solution is to make use of anatomical priors to allow for the traversal of GM structures in cases where connections are known to pass through (e.g. allowing connections to pass through GPi when connecting GPe and STN). Such a solution has been previously implemented for cortico-cortical connectivity in the White Matter Query Language, where predefined regions (inclusion and exclusion) and endpoint ROIs are used to identify connections of interest (Wassermann et al., 2016). To implement this for the subcortical connectome would require detailed curation of anatomical knowledge to identify the necessary inclusion and exclusion wayward ROIs required in addition to the terminal regions. Unfortunately, even without explicit exclusion of wayward GM ROIs, the ability for tracts to pass through GM will be challenging due to the reduced anisotropy in GM (e.g. in the pallidum).

In a similar manner, the inclusion of WM priors as wayward ROIs may improve anatomical accuracy. With a data-driven approach to identifying the subcortical connectome, we had observed the presence of major tracts (e.g. CST), spurious streamlines, and in some instances, multiple trajectories between two subcortical structures. By using a WM prior, trajectories from major tracts and spurious streamlines could be filtered, while individual trajectories can be isolated. In a previous study of tractography reproducibility, a suggestion was made to include the use of anatomical priors as guidance to improve identification of connectivity (Maier-Hein et al., 2017). One such possibility is to leverage the segmentations of subcortical connections surrounding the zona incerta that have been previously identified with high resolution, *in vivo* anatomical MRI (Lau et al., 2020) to help differentiate observed connections from a data-driven approach. Additionally, drawing anatomical knowledge from NHP and post-mortem studies can help to establish priors that can improve anatomical accuracy by minimizing the number of false positive connections and help to discern trajectories. However, optimizing the use of anatomical priors remains an open challenge even for well understood tracts like the CST (Rheault et al., 2020; Schilling et al., 2021). Ultimately, moving forward, our described framework would allow for the evaluation of different implementation choices and their impact on both identification and reliability of mapping the subcortical connectome.

### 4.5 Limitations

Several limitations are worth noting beyond those related to choices made in the implementation of the trajectory algorithm (see previous section). Validation of tractography identified connections *in vivo* is a known challenge, given the limited ability to compare to ground truth trajectories, which have been conventionally identified using tract-tracing in experimental animals. While the most accurate comparisons would be performed between tract-tracing and tractography on the same brain, this is not feasible in humans. Fortunately, connections between regions are highly similar across different primates (Grisot et al., 2021). In the current study, we limited our investigation to known connections between ROIs of the basal ganglia and the thalamus in order to compare our observations with previously described trajectories. As a result, we did not explore the complete network circuitry to other regions of the brain (e.g. brainstem, cerebellum, cortex, etc). Some of these unexplored regions contain important nodes, such as connectivity with the hypothalamus (Haber et al., 1993) and pedunculopontine nucleus of the brainstem (DeLong and Wichmann, 2007). Other connections of interest, including between the sensory thalamus (e.g. medial and lateral geniculate nuclei) and the striatum, have been previously examined in experimental animals (Takada et al., 1985). Future work should explore the network circuitry more comprehensively, which should be increasingly feasible with increasing availability of brain atlases.

## 5. Conclusion

In this study, we demonstrated that identifying the subcortical connectome using a data-driven probabilistic approach with *in vivo* tractography was both feasible and reliable, with a particular focus on the assessment of known connections that have been previously described. Quantitative evaluation of the subcortical connections demonstrated similar tract densities and overlap comparable to what has been shown in existing studies focused on cortico-cortical and cortico-subcortical networks. Performing this assessment also highlighted areas requiring improvement to address the challenges of tractography in the subcortex. The methods used in this study can serve as a framework for evaluating the impact of modifications to the tractography workflow, with the goal of increasingly accurate and reliable mapping of the subcortical connectome.

## Data Availability

The data used in this study are available as part of the publicly available Human Connectome Project S1200 release (https://humanconnectome.org/study/hcp-young-adult). Analysis was performed with JupyterLab using Python (version 3.8) and the code is available at https://github.com/kaitj/hcp_subcortical_repro.

## Ethics Statement

All participants provided their written informed consent. The use of a publicly available, open dataset was approved by the University of Western Ontario Health Sciences Research Ethics Board (UWO HSREB) under protocol #108456 (PI: Khan).

## Acknowledgements

JK was supported by the postgraduate scholarship from the National Sciences and Engineering Research Council (NSERC). RH was supported by a BrainsCAN postdoctoral fellowship. AK was supported by the Canada Research Chairs program #950-231964, NSERC Discovery Grant #6639, and Canada Foundation for Innovation (CFI) John R. Evans Leaders Fund Project #37427, the Canada First Research Excellence Fund, and Brain Canada. JL was supported by research start-up funding through the Department of Clinical Neurological Sciences at University of Western Ontario. This research was enabled in part by the support provided by Compute Ontario (www.computeontario.ca) and Compute Canada (www.computecanada.ca). Data was provided in part by the Human Connectome Project, WU-Minn Consortium (Principal Investigators: David Van Essen and Kamil Ugurbil; 1U54MH091657) funded by the 16 NIH Institutes and Centers that support the NIH Blueprint for Neuroscience Research; and by the McDonnell Center for Systems Neuroscience at Washington University.

## Supplementary Materials - Results from HCP Unrelated dataset

### S1.1 Networks of the subcortical connectome

Known connections within the networks of interest (i.e. motor, associative, and limbic) were examined for the HCP unrelated dataset. Fig. 2 demonstrated the connectivity and the associated TDs for the assessed motor network (see Supplementary Fig. 2A and Supplementary Fig. 2B for associative and limbic network). Using the same TD threshold of 6.5 AFD previously determined, connectivity was observed to be similar to both sessions of the test-retest dataset. 78% (14 out of 18) of the known connections of the motor network met the previously determined threshold, while 100%% and 71% (10 out of 14) connections of the associative and limbic networks satisfied the threshold respectively. Moderate density between subcortical structures of interest were observed except with certain thalamic nuclei as before. The same subcortical connections which had previously failed to meet the tract density threshold also failed to meet the threshold for the HCP unrelated dataset. Visual inspection of connectivity noted identical observations as with the test-retest dataset with shorter, more direct connections between basal ganglia structures, and longer connections with a more curved trajectory between the basal ganglia and thalamus.

### S1.2 Reliability of the HCP unrelated dataset

Connectivity matrices were first created for subjects of the HCP unrelated dataset, before an average TD matrix across subjects was computed and examined (Fig. S1A). Furthermore, a box plot of the TD exhibited in each hemisphere was created and overlaid by a swarmplot of individual TDs between subcortical structures of interest (Fig. S1B). A visual comparison with similar plots created for the test-retest dataset suggested comparable connectivity being demonstrated in the unrelated subjects. A Pearson’s correlation was performed between the average TD of the unrelated dataset and the average TD for the test and retest sessions respectively, where a Pearson’s correlation coefficient of 0.99 was exhibited against both test (p < 0.05) and retest sessions (p < 0.05).

**Figure S1.**
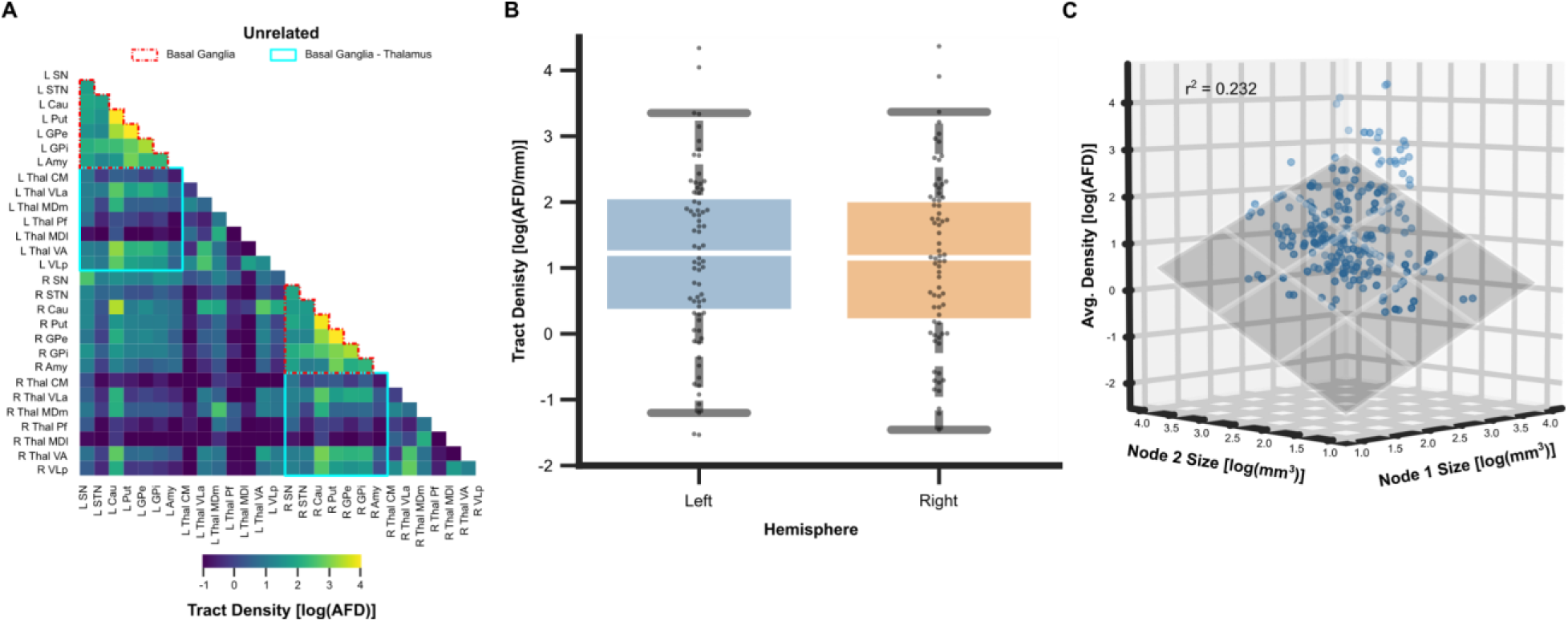
(A) Log-transformed average tract density of the HCP unrelated subjects are shown in a connectivity matrix, visualizing connectivity between subcortical structures of known subcortical networks. (B) The tract density, exhibited in a box plot and overlaid with a swarmplot to exhibit individual TDs, is separated by hemispheric connectivity. The middle line of the boxplot marks the median metric, while whiskers define the maximum and minimum values of each metric, excluding outliers. Average tract densities similar to those previously exhibited in both HCP test and retest sessions. (C) A 3-dimensional scatterplot is shown, observing the relationship between the average tract density of connections and the volume of the terminal nodes. A similar relationship to that from the test-retest dataset was observed.

The tract density (TD) was evaluated against the size of the two connecting subcortical structures for the unrelated dataset. A positive linear relationship was observed (Fig S1C; r^2^ = 0.232, p < 0.05), with an increase in average TD associated with an increase in size of one or both subcortical structures, demonstrating a positive linear relationship when an ordinary least squares regression was performed.

**Supplementary Table 1.**
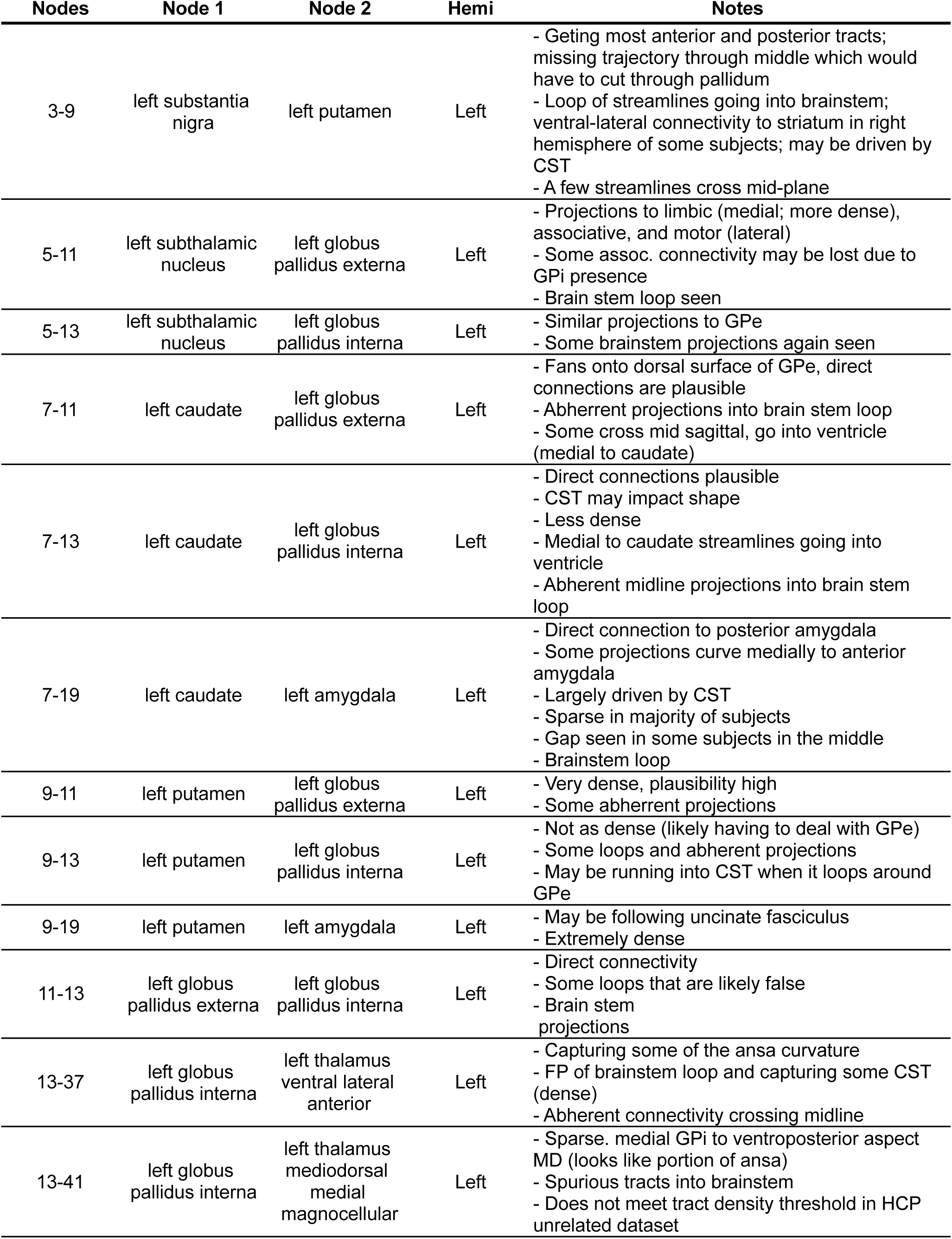
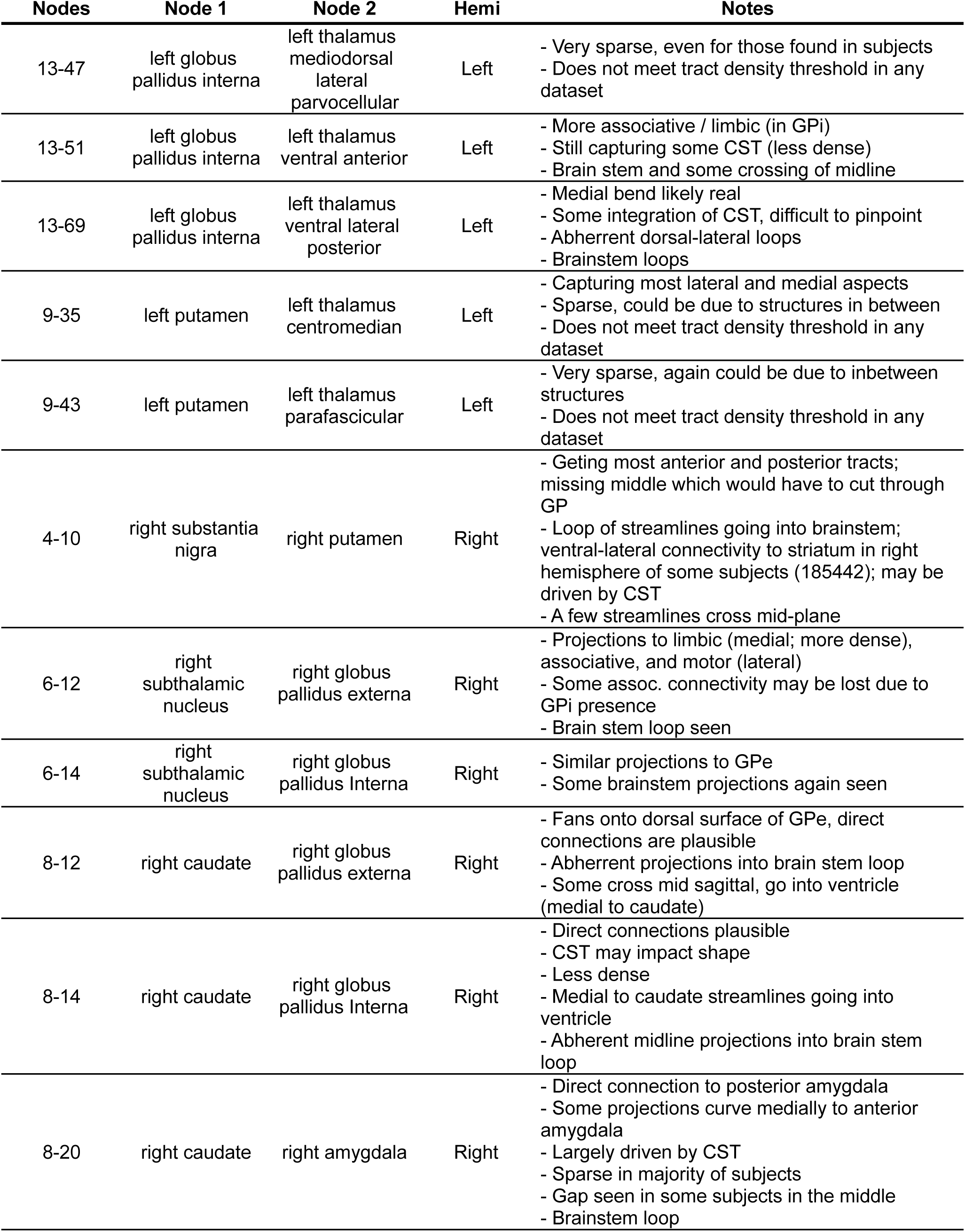
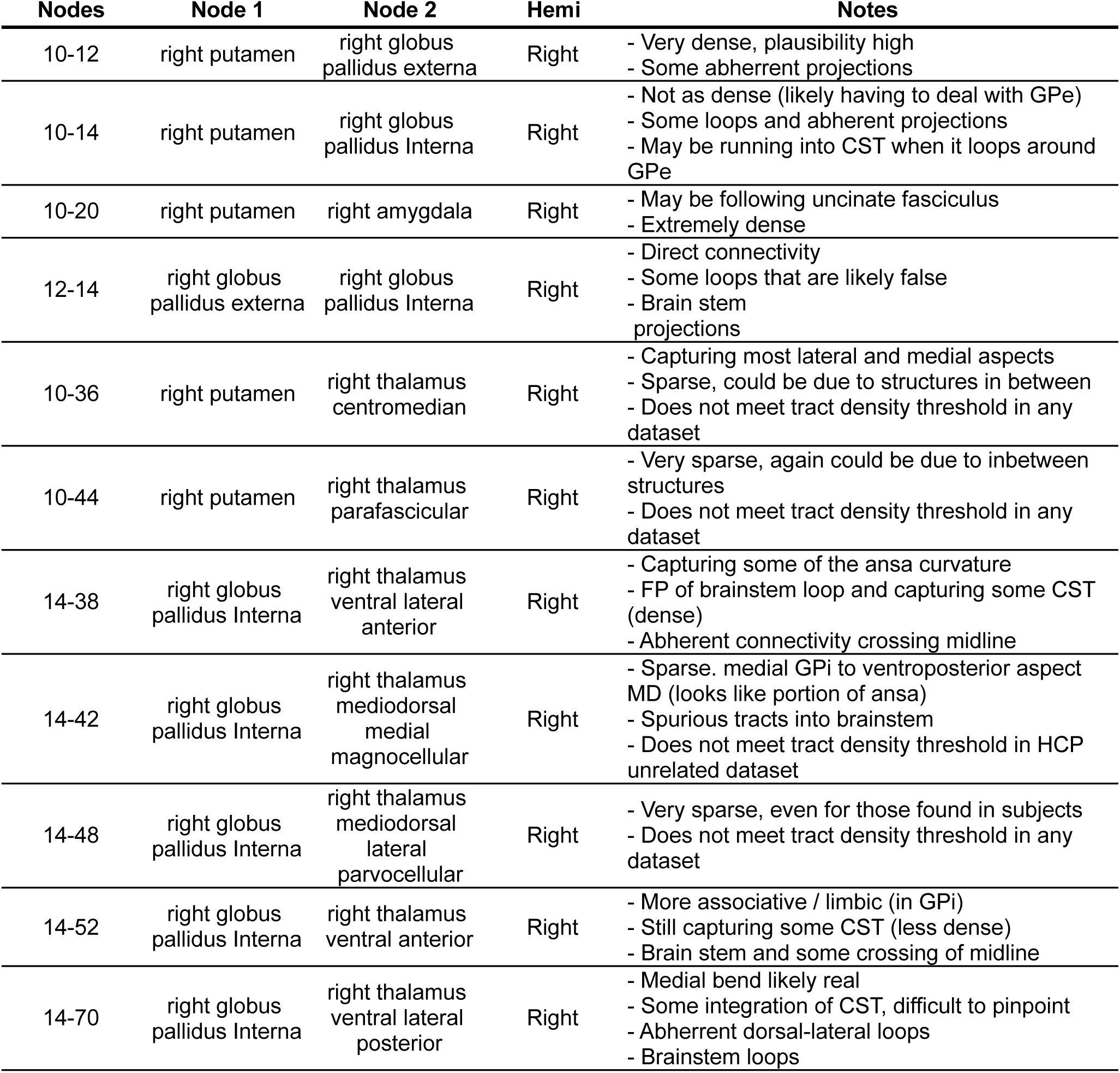
Visual observations of tract trajectories between subcortical structures

| **Nodes** | **Node 1** | **Node 2** | **Hemi** | **Notes** |
| --- | --- | --- | --- | --- |
| 3-9 | left substantia nigra | left putamen | Left | - Getting most anterior and posterior tracts; missing trajectory through middle which would have to cut through pallidum<br>- Loop of streamlines going into brainstem; ventral-lateral connectivity to striatum in right hemisphere of some subjects; may be driven by CST<br>- A few streamlines cross mid-plane |
| 5-11 | left subthalamic nucleus | left globus pallidus externa | Left | - Projections to limbic (medial; more dense), associative, and motor (lateral)<br>- Some assoc. connectivity may be lost due to GPi presence<br>- Brain stem loop seen |
| 5-13 | left subthalamic nucleus | left globus pallidus interna | Left | - Similar projections to GPe<br>- Some brainstem projections again seen |
| 7-11 | left caudate | left globus pallidus externa | Left | - Fans onto dorsal surface of GPe, direct connections are plausible<br>- Abherrent projections into brain stem loop<br>- Some cross mid sagittal, go into ventricle (medial to caudate) |
| 7-13 | left caudate | left globus pallidus interna | Left | - Direct connections plausible<br>- CST may impact shape<br>- Less dense<br>- Medial to caudate streamlines going into ventricle<br>- Abherent midline projections into brain stem loop |
| 7-19 | left caudate | left amygdala | Left | - Direct connection to posterior amygdala<br>- Some projections curve medially to anterior amygdala<br>- Largely driven by CST<br>- Sparse in majority of subjects<br>- Gap seen in some subjects in the middle<br>- Brainstem loop |
| 9-11 | left putamen | left globus pallidus externa | Left | - Very dense, plausibility high<br>- Some abherrent projections |
| 9-13 | left putamen | left globus pallidus interna | Left | - Not as dense (likely having to deal with GPe)<br>- Some loops and abherent projections<br>- May be running into CST when it loops around GPe |
| 9-19 | left putamen | left amygdala | Left | - May be following uncinate fasciculus<br>- Extremely dense |
| 11-13 | left globus pallidus externa | left globus pallidus interna | Left | - Direct connectivity<br>- Some loops that are likely false<br>- Brain stem projections |
| 13-37 | left globus pallidus interna | left thalamus ventral lateral anterior | Left | - Capturing some of the ansa curvature<br>- FP of brainstem loop and capturing some CST (dense)<br>- Abherent connectivity crossing midline |
| 13-41 | left globus pallidus interna | left thalamus mediodorsal medial magnocellular | Left | - Sparse. medial GPi to ventroposterior aspect MD (looks like portion of ansa)<br>- Spurious tracts into brainstem<br>- Does not meet tract density threshold in HCP unrelated dataset |

| **Nodes** | **Node 1** | **Node 2** | **Hemi** | **Notes** |
| --- | --- | --- | --- | --- |
| 13-47 | left globus pallidus interna | left thalamus mediodorsal lateral parvocellular | Left | - Very sparse, even for those found in subjects<br>- Does not meet tract density threshold in any dataset |
| 13-51 | left globus pallidus interna | left thalamus ventral anterior | Left | - More associative / limbic (in GPi)<br>- Still capturing some CST (less dense)<br>- Brain stem and some crossing of midline |
| 13-69 | left globus pallidus interna | left thalamus ventral lateral posterior | Left | - Medial bend likely real<br>- Some integration of CST, difficult to pinpoint<br>- Abherrent dorsal-lateral loops<br>- Brainstem loops |
| 9-35 | left putamen | left thalamus centromedian | Left | - Capturing most lateral and medial aspects<br>- Sparse, could be due to structures in between<br>- Does not meet tract density threshold in any dataset |
| 9-43 | left putamen | left thalamus parafascicular | Left | - Very sparse, again could be due to inbetween structures<br>- Does not meet tract density threshold in any dataset |
| 4-10 | right substantia nigra | right putamen | Right | - Getting most anterior and posterior tracts; missing middle which would have to cut through GP<br>- Loop of streamlines going into brainstem; ventral-lateral connectivity to striatum in right hemisphere of some subjects (185442); may be driven by CST<br>- A few streamlines cross mid-plane |
| 6-12 | right subthalamic nucleus | right globus pallidus externa | Right | - Projections to limbic (medial; more dense), associative, and motor (lateral)<br>- Some assoc. connectivity may be lost due to GPi presence<br>- Brain stem loop seen |
| 6-14 | right subthalamic nucleus | right globus pallidus Interna | Right | - Similar projections to GPe<br>- Some brainstem projections again seen |
| 8-12 | right caudate | right globus pallidus externa | Right | - Fans onto dorsal surface of GPe, direct connections are plausible<br>- Abherrent projections into brain stem loop<br>- Some cross mid sagittal, go into ventricle (medial to caudate) |
| 8-14 | right caudate | right globus pallidus Interna | Right | - Direct connections plausible<br>- CST may impact shape<br>- Less dense<br>- Medial to caudate streamlines going into ventricle<br>- Abherent midline projections into brain stem loop |
| 8-20 | right caudate | right amygdala | Right | - Direct connection to posterior amygdala<br>- Some projections curve medially to anterior amygdala<br>- Largely driven by CST<br>- Sparse in majority of subjects<br>- Gap seen in some subjects in the middle<br>- Brainstem loop |

| **Nodes** | **Node 1** | **Node 2** | **Hemi** | **Notes** |
| --- | --- | --- | --- | --- |
| 10-12 | right putamen | right globus pallidus externa | Right | - Very dense, plausibility high<br>- Some abherrent projections |
| 10-14 | right putamen | right globus pallidus Interna | Right | - Not as dense (likely having to deal with GPe)<br>- Some loops and abherrent projections<br>- May be running into CST when it loops around GPe |
| 10-20 | right putamen | right amygdala | Right | - May be following uncinate fasciculus<br>- Extremely dense |
| 12-14 | right globus pallidus externa | right globus pallidus Interna | Right | - Direct connectivity<br>- Some loops that are likely false<br>- Brain stem projections |
| 10-36 | right putamen | right thalamus centromedian | Right | - Capturing most lateral and medial aspects<br>- Sparse, could be due to structures in between<br>- Does not meet tract density threshold in any dataset |
| 10-44 | right putamen | right thalamus parafascicular | Right | - Very sparse, again could be due to inbetween structures<br>- Does not meet tract density threshold in any dataset |
| 14-38 | right globus pallidus Interna | right thalamus ventral lateral anterior | Right | - Capturing some of the ansa curvature<br>- FP of brainstem loop and capturing some CST (dense)<br>- Abherrent connectivity crossing midline |
| 14-42 | right globus pallidus Interna | right thalamus mediodorsal medial magnocellular | Right | - Sparse. medial GPI to ventroposterior aspect MD (looks like portion of ansa)<br>- Spurious tracts into brainstem<br>- Does not meet tract density threshold in HCP unrelated dataset |
| 14-48 | right globus pallidus Interna | right thalamus mediodorsal lateral parvocellular | Right | - Very sparse, even for those found in subjects<br>- Does not meet tract density threshold in any dataset |
| 14-52 | right globus pallidus Interna | right thalamus ventral anterior | Right | - More associative / limbic (in GPI)<br>- Still capturing some CST (less dense)<br>- Brain stem and some crossing of midline |
| 14-70 | right globus pallidus Interna | right thalamus ventral lateral posterior | Right | - Medial bend likely real<br>- Some integration of CST, difficult to pinpoint<br>- Abherrent dorsal-lateral loops<br>- Brainstem loops |

---

**Supplementary Figure 1.**
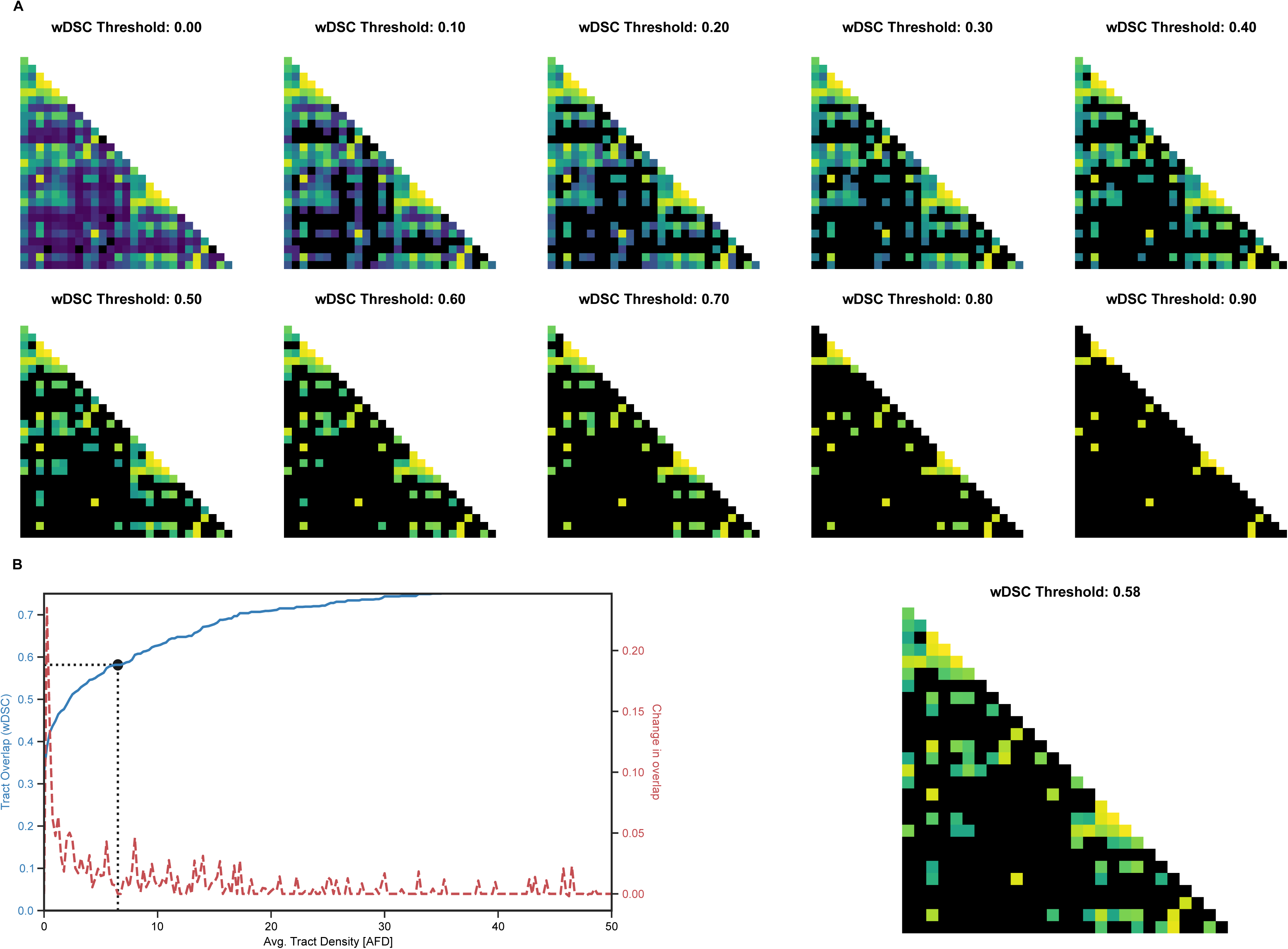
(A) Example of connectivity matrices at difference spatial overlap (wDSC) thresholds. Connections unable to meet the thresholds are discarded and are shown in the connectivity matrices as black boxes. (B) Plot demonstrating how the threshold is chosen by sweeping through tract densities and identifying both the tract overlap and the change in overlap between consecutive thresholds (left). Connectivity matrix at the selected threshold of wDSC = 0.58 (right).

**Supplementary Figure 2.**
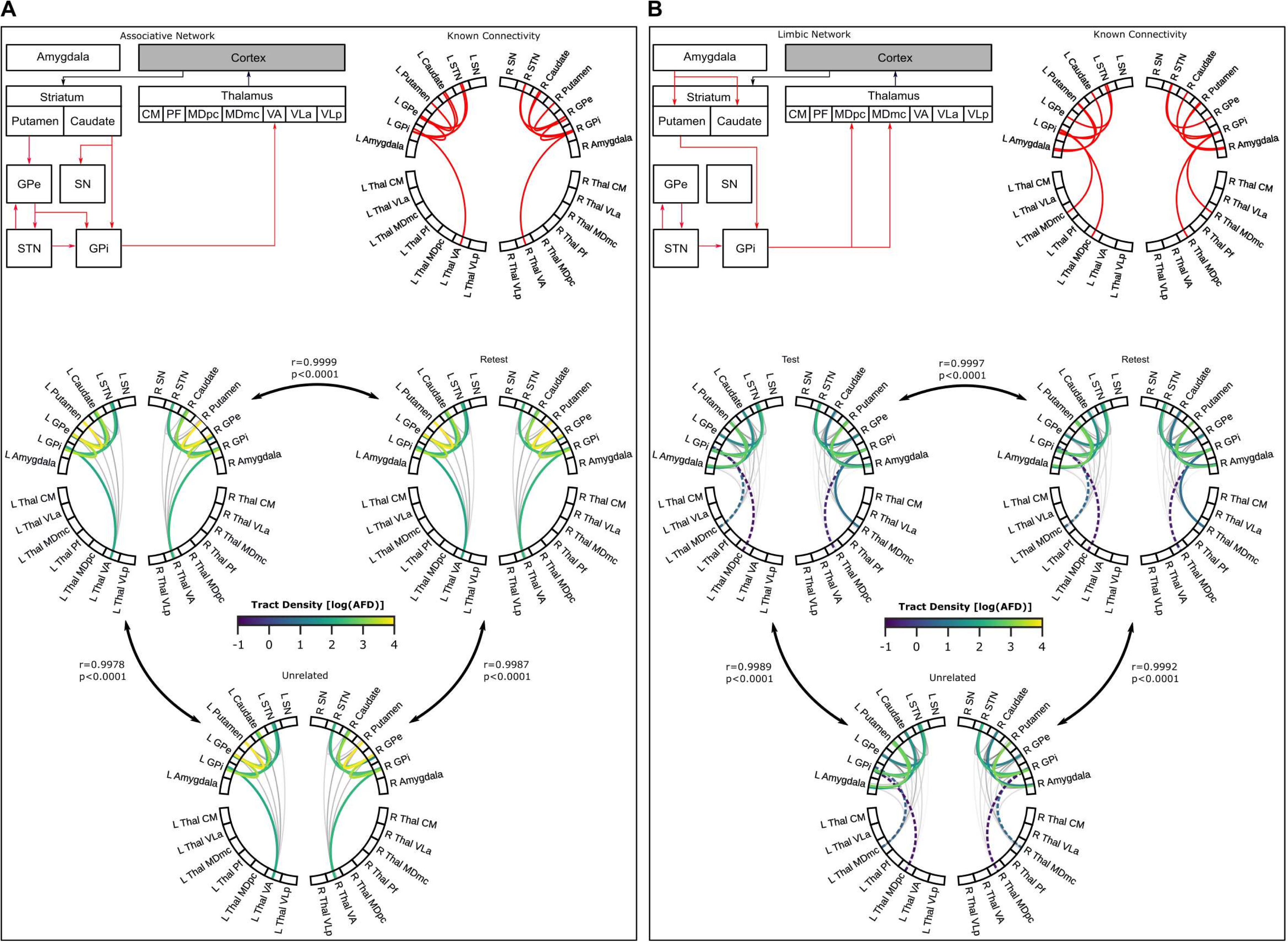
Known subcortical connections (in red) of the (A) associative network and (B) limbic network. For each network, connectivity identified from literature was depicted in a diagram (top-left) and chord plot (top-right), while chord plots exhibit the average log-transformed tract densities for test-retest (middle-left, middle-right) and unrelated (bottom) datasets of the Human Connectome Project. Dashed lines represent connections that did not meet the selected tract density threshold. Pearson correlations between datasets are shown next to the comparison indicators.

**Supplementary Figure 3.**
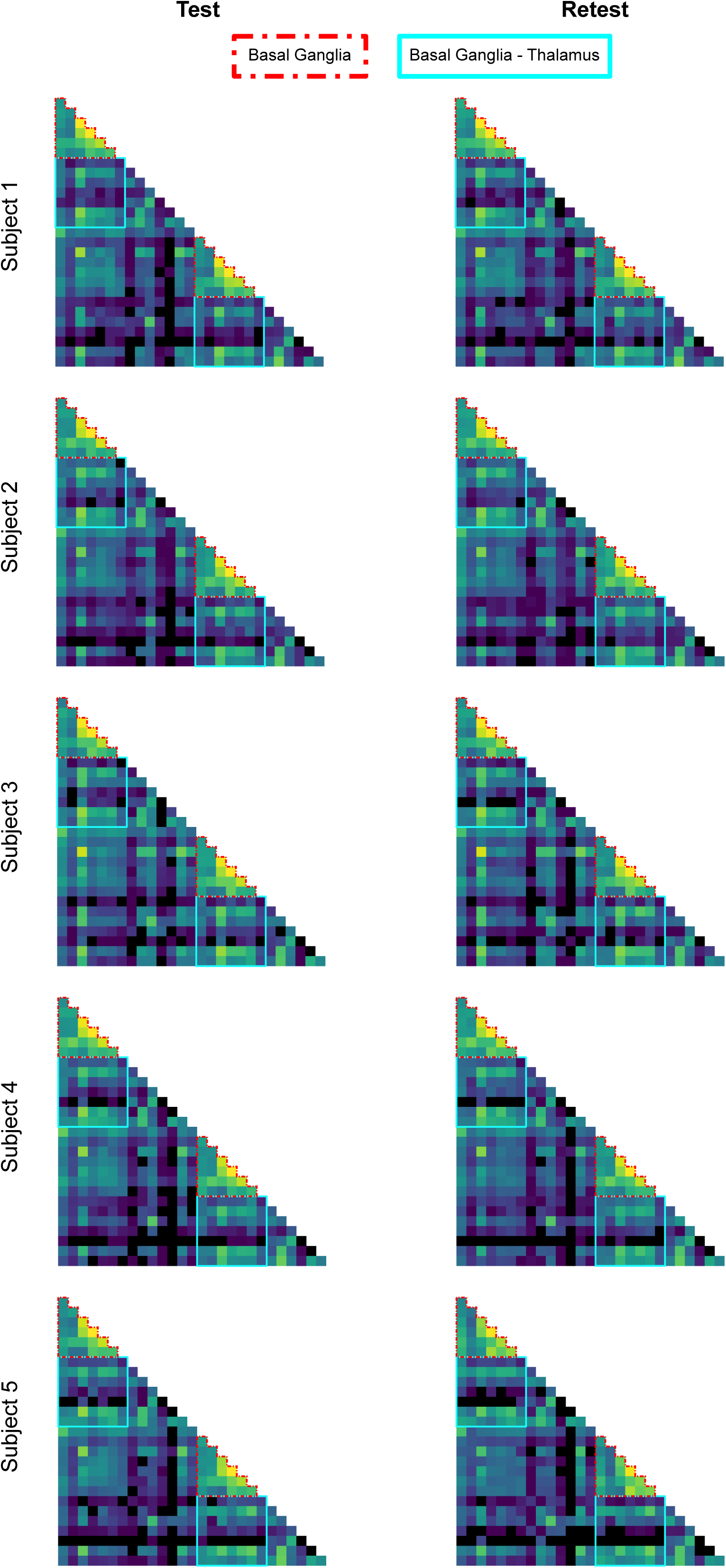
Examples of connectivity matrices exhibiting tract density are shown for 5 different subjects, with connectivity from the test session on the left and retest session on the right. Outlined boxes highlight the connectivity of the basal ganglia (red) and between the basal ganglia and thalamus (cyan).

**Supplementary Figure 4.**
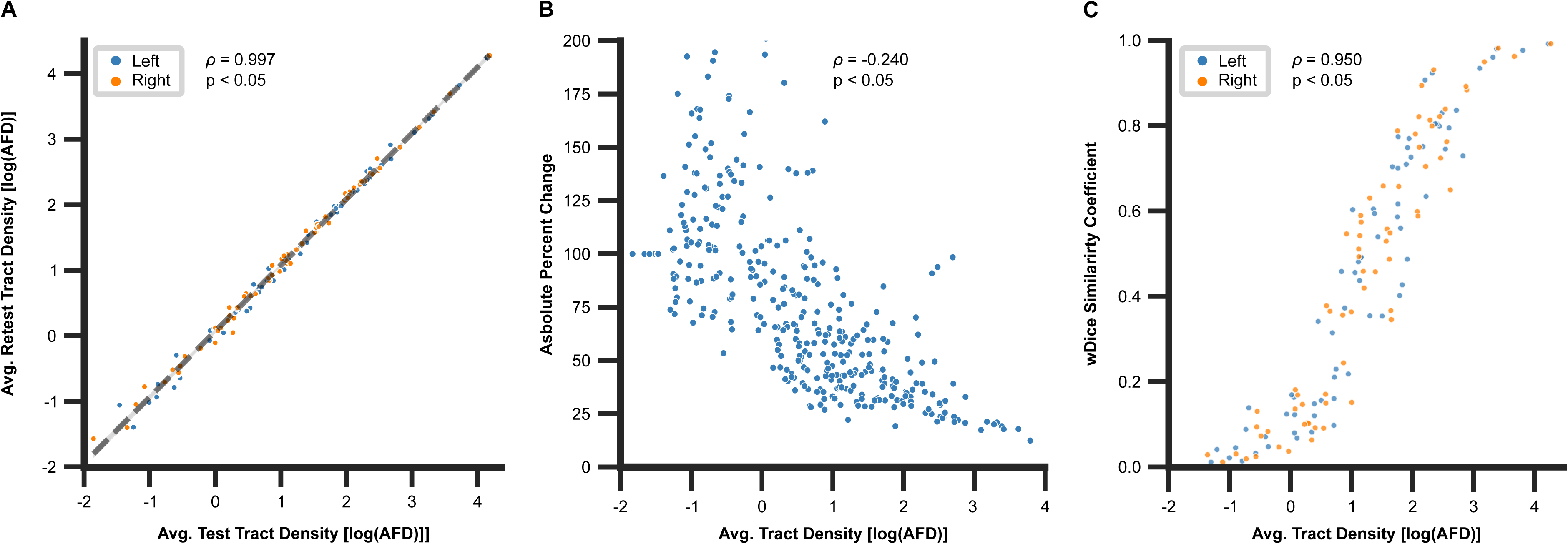
Examples of connectivity matrices exhibiting tract density are shown for 5 different subjects, with connectivity from the test session on the left and retest session on the right. Outlined boxes highlight the connectivity of the basal ganglia (red) and between the basal ganglia and thalamus (cyan).

1 https://humanconnectome.org/study/hcp-young-adult/document/1200-subjects-data-release

